# Reproductive isolation due to divergent ecological selection is accompanied by vast genomic instability in experimentally evolved yeast populations

**DOI:** 10.1101/2025.01.28.635359

**Authors:** Devin P. Bendixsen, Ciaran Gilchrist, Chloé Haberkorn, Karl Persson, Cecilia Geijer, Jonas Warringer, Rike Stelkens

## Abstract

Populations evolving independently in divergent environments accumulate genetic differences and potentially evolve reproductive isolation as a by-product of divergence. The speed and mechanisms underlying this process are difficult to investigate because we rarely get the opportunity to witness them in natural settings, and histories of selection and gene flow between populations are often unknown. Here, we experimentally evolved yeast for 1000 generations of evolution in both divergent and parallel environments. At regular time points during experimental evolution, we made crosses between parallel- and divergent-evolving populations to measure postzygotic reproductive isolation (gamete viability). We used whole genome population sequencing to determine the mutational load, the number and types of structural variation, and other genomic features of the parent, F1, and F2 intraspecific hybrids. We found evidence for large scale phenotypic and genome-wide differentiation in response to divergent laboratory selection. Divergent-selected populations produced hybrids with reduced gamete viability - a classic signature of postzygotic reproductive isolation in the form of hybrid breakdown. Parallel-selected populations on the other hand remained more reproductively compatible (with exceptions). We found that F2 hybrid genomes contained vast genomic instability, i.e., new structural variants (especially insertions, deletions, and interchromosomal translocations) that were not observed in parent and F1 genomes, which is likely a result of chromosome missegregation and recombination errors in hybrid meiosis. Our results provide phenotypic and genomic evidence that partial reproductive isolation evolved due to adaptation to divergent environments, consistent with predictions of ecological speciation theory.

## Introduction

Ecological speciation implies a process by which adaptation to different ecological niches, in allopatry or sympatry, leads to the evolution of reproductive isolation between populations (Mayr 1947; Rice and Hostert 1993; Schluter 2001; McKinnon et al. 2004; Nosil 2012). Long-term monitoring of natural populations has been valuable in linking ecological traits with the emergence of reproductive barriers (e.g. (Filchak et al. 2000; Rundle et al. 2000; Grant and Grant 2003; Funk et al. 2006; Sawada et al. 2022)), providing convincing arguments for ecological factors to play a role in driving population divergence. However, ecological speciation remains difficult to demonstrate because it usually proceeds over extended evolutionary times (Hendry et al. 2007), trait divergence is often cryptic (Struck et al. 2018; Sparks et al. 2022), and the evolutionary histories of natural populations are challenging to reconstruct. It is usually unknown whether divergent selection was present, how long it lasted, and what the timing and amount of gene flow was between populations. Thus, it often remains unclear, especially between allopatric species pairs occupying different niches, whether divergent, ecology-based selection was the driving force underlying their divergence, eventually leading to speciation (Anderson and Weir 2022; Irestedt et al.; Müller et al.). Due to these constraints, studies have often relied on combinatorial approaches, using i) data on niche divergence in traits related to habitat use, such as resource utilization or foraging behavior (e.g. (Hatfield and Schluter 1999; Nosil and Crespi 2006; Munar-Delgado et al. 2024), ii) data on reproductive barriers, such as differences in mating behaviors or flowering times reducing gene flow between populations adapted to different ecological niches (e.g. (Dieckmann et al. 2004; Villa et al. 2019)), and iii) genetic data, detecting signatures of selection and divergence in genetic regions associated with ecological adaptation and reproductive isolation (e.g. (Nosil and Schluter 2011; Härer et al. 2021; Louder et al. 2024)).

Microbial experimental evolution provides a powerful tool for investigating questions in evolutionary biology. It has been used to describe the dynamics and genetic mechanisms underlying adaptation to stable (Lenski et al. 1991; Good et al. 2017; Johnson and Desai 2022), heterogenous (Rainey and Travisano 1998), or changing environments (Bell and Gonzalez 2011; Gorter et al. 2016; Gorter et al. 2017) and to test theoretical predictions about adaptation rates in sexual versus asexual populations (McDonald et al. 2016; Kosheleva and Desai 2018; Leu et al. 2020). Experimental evolution with microbes can also provide insights on fundamental questions in speciation, especially about the early stages of divergence, i.e. why, how and when lineages split (Stelkens 2024). Microbial genomes are often small and generation times are short, allowing for evolve-and-resequencing approaches to capture speciation and its underlying genetic basis in action. For instance, experimental evolution with a virus (lambda bacteriophage) and its bacterial host (*Escherichia coli*) has demonstrated that divergent ecological selection caused the phages to split into two genetically distinct lineages with different host preferences, and that barriers to gene flow formed as a consequence of ecological specialization, which resulted in reproductive isolation (Meyer et al. 2016). Experiments with yeast and other fungi have shown that divergent selection can lead to phenotypic divergence and rapid ecological differentiation (Dettman et al. 2007; Dettman et al. 2008; Ament-Velásquez et al. 2022), even under high rates of gene flow (Tusso et al. 2021). But generally, there is a lack of studies that combine both organismal and genetic approaches to measure reproductive isolation (Westram et al. 2022). Work investigating the emergence and dynamics of reproductive isolation using repeated measures of hybrid viability across generational time scales, from the beginning to later stages of parental divergence under controlled environmental conditions, could significantly advance our understanding of what can drive the evolution of reproductive isolation in the early stages of speciation.

Here, we used experimental evolution with the budding yeast *Saccharomyces cerevisiae* and propagated replicate, asexually reproducing populations in both different and parallel (same) environments for 1000 mitotic generations (**Figure 1**). At regular time points of population divergence, we generated sexual F1 and F2 (hybrid) crosses between and within environments, to test whether reproductive isolation evolved and increased with time. We evolved populations in four distinct selective environments and profiled parental phenotypes in 90 different environments to test for phenotypic divergence over time. We then used whole genome sequencing on parental, F1, and F2 hybrid populations to measure their mutational load and other measures of genomic stability. By the endpoint of experimental evolution, we found that divergent-selected populations were up to 50% reproductively isolated due reduced hybrid gamete viability, while most crosses made between parallel-selected populations remained genetically compatible. We detected vast structural variation in the F2 hybrid genomes, including up to three times more deletions, insertions, inversions, tandem duplications, and interchromosomal translocations than in the diploid parental and F1 hybrid genomes. Consistent with predictions of ecological speciation theory, our results suggest that divergent adaptation has led to partial reproductive isolation, accompanied by severe genomic instability in the hybrids. This instability may be the result of structural genetic differences accumulating in parental populations causing recombination errors and chromosome missegregation during hybrid meiosis, but further work is needed to test if this genetic mechanism underlies the evolution of reproductive isolation in our system.

**Figure 1:**
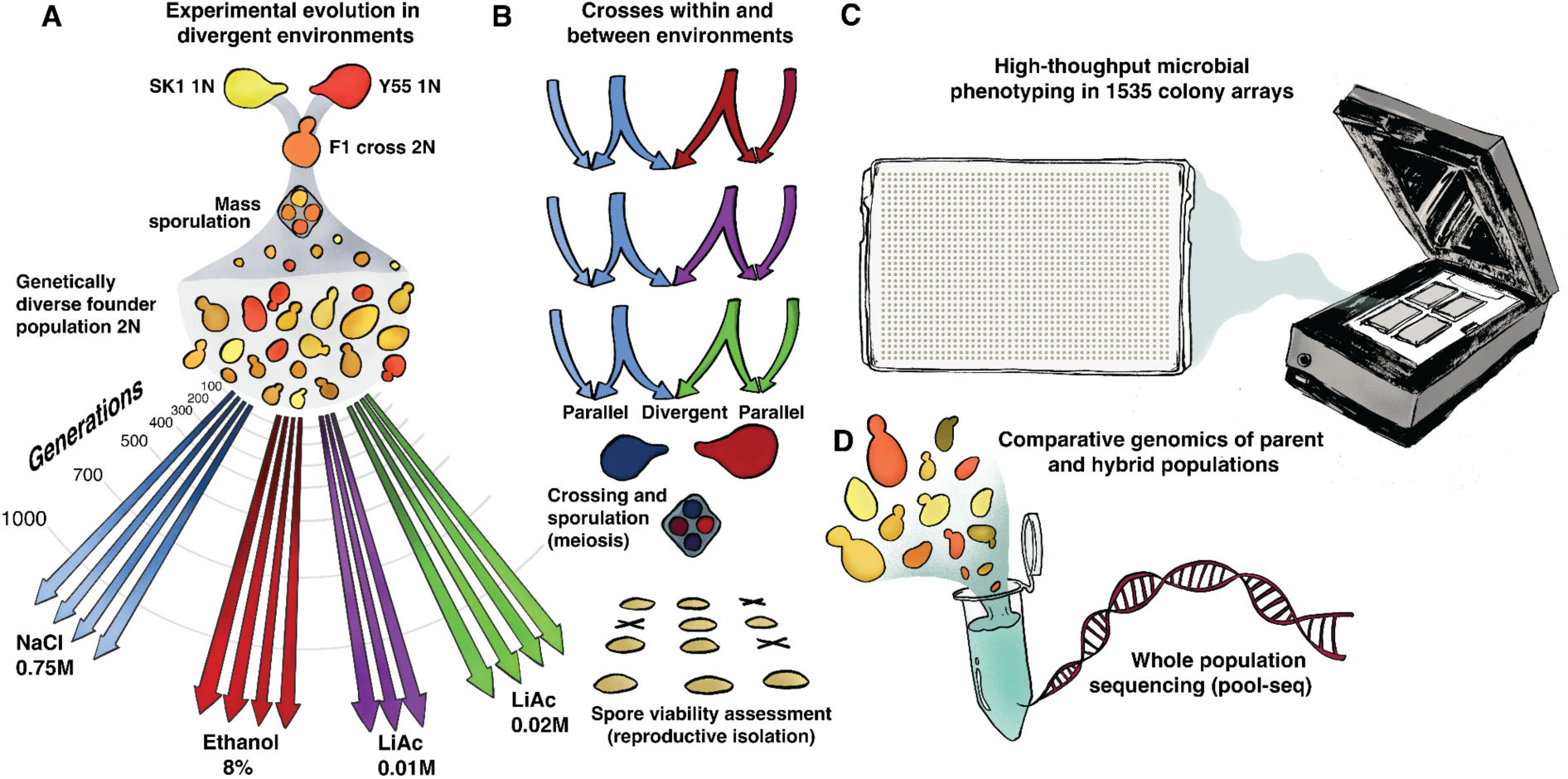
Overview of experimental approaches. **A) Experimental Evolution**: A founder population was generated by the mating of two *S. cerevisiae* strains (SK1 and Y55), followed by mass sporulation (meiosis). This genetically diverse, diploid founder was split into four different environments, containing growth medium with either 0.75M NaCl, 8% EtOH, 0.01M lithium acetate, or 0.02M lithium acetate, and evolved for up to 1000 asexual generations by serial transfer, in three or four independent biological replicates per environment (Ament-Velásquez et al. 2022). 2N indicates diploid, 1N indicates haploid genotypes. **B) Crossing design**: At seven points of parental divergence (indicated by the curved grey lines in A), sexual hybrid crosses were made between NaCl-populations and populations from all other environments (with replication). Control crosses were made between parallel-evolved, replicate populations from the same environment. Crosses from five time points were then sporulated and the viability of the haploid F2 offspring was assessed after tetrad dissection. **C) High throughput phenotyping:** To assess phenotypic divergence, the growth of parental populations sampled from five time points of experimental evolution was measured in 90 non-selection environments using colony image scans. **D) Comparative genomics** using pool sequencing of parental, F1, and F2 populations sampled at seven time points of experimental evolution.

## Materials and Methods

### Founder populations

We generated large diploid, and genetically diverse, founder populations of yeast through mating and mass sporulation of two related *S. cerevisiae* strains (SK1 and Y55; Figure 1) as previously described (Ament-Velásquez et al. 2022), for which high quality reference genomes are available (Peter et al. 2018; Bendixsen, Gettle, et al. 2021). Briefly, SK1 and Y55 were genetically modified to introduce auxotrophies, which were used as selective markers to exclude non-mated haploids (non-hybrids). From these strains, two founding populations were generated that only differed in the selective markers they carried. These selective markers allowed us to mass-mate the NaCl-evolved populations with populations evolving in the other three environments, while safely removing all non-hybrid parental genotypes, using a selective growth medium. In the synthetic complete growth medium we used for experimental evolution, these auxotrophic markers have no detectable fitness effect as all essential amino acids are provided. The founders of populations evolving in NaCl (called “N founder” below) are the diploid recombined offspring of a cross between SK1 (MATα ho:KanMX ura3::ClonNat) and Y55 (MATa ho leu2::HygMX ura3::ClonNat). Founders of the populations evolving in ethanol and two different concentrations of lithium acetate (called “LE founder” below), are the diploid recombined offspring of a cross between SK1 (MATα ho:KanMX lys2::ClonNat) and Y55 (MATa ho leu2::HygMX lys2::ClonNat). Despite having different selective markers, it is important to note that both founders maintain the same strain background and differ only in their fitness-neutral auxotrophic markers.

### Experimental evolution of parental populations

From these founders, replicate populations (from here on ‘parentaĺ populations) were experimentally evolved in four divergent environments, each with four replicates (considered parallel-evolved) for up to 1000 asexual generations (**Figure 1**). The number of generations was estimated by counting cells after 48h growth in selective media in the highest stress environment (0.75M NaCl, see Ament-Velásquez et al. 2022). This approach is conservative because it likely underestimates the number of generations in the other, less stressful environments. All evolution environments contained synthetic complete (SC) medium (2% glucose, 0.67% bacto-yeast nitrogen base without amino acids, 0.00079% Formedium CSM powder (product code DCS0019), made up to 1L with sterilised deionised water) plus one of the following substances to induce stress: 0.75M sodium chloride (from here on NaCl), 8% ethanol (C₂H₆O, from here on EtOH), and two different concentrations of lithium acetate (C_2_H_3_LiO_2_, from here on LiAc0.01M and LiAc0.02M). The correct volume of EtOH was added to the test tube just before inoculation to limit evaporation. We have previously demonstrated that these four media significantly differ in how much they reduce the fitness of populations, and we have provided phenotypic and genetic evidence that populations become differentially adapted to these environments (Ament-Velásquez et al. 2022).

Three to four replicate populations were evolved in separate test tubes containing 5mL of SC media with one of the four substances. Then, we serially transferred 10µL of yeast culture every 48 hours to fresh media (approximately seven generations, estimated by counting cells after 48 hours). Every five transfers (10 days, i.e. ∼35 generations), a 900µL sample of each population was frozen in 44% glycerol at −70°C for later use. The experiment was run for a total of ∼300 days. After some cross-replicate contamination was detected using sequencing data (Ament-Velásquez et al. 2022), we excluded two NaCl-evolved replicate populations (R1 and R4) at 1000 generations, all LiAc0.01M and LiAc0.02M-evolved populations at 1000 generations, and one LiAc0.01M-evolved population (R3) at all generations. No crosses were generated from these contaminated populations. Retained populations showed no signs of cross-contamination in terms of shared single nucleotide variants.

### Hybridization at increasing timepoints of parental divergence

We generated hybrid crosses at 100, 300, 500, 700, and 1000 generations of parental divergence, between all four replicate NaCl-populations (excluding R1 and R4 at 1000 generations due to contamination) and up to four replicate populations from each of the three other environments (LiAc 0.01M (3 replicates), LiAc 0.02M (4 replicates) and EtOH (4 replicates); **Figure 1**). Due to the selectable markers present in the parents, NaCl-populations could be mass-crossed with the other three populations while safely removing all non-hybrid parental genotypes using a selective medium (described below). First, we mass-sporulated the diploid parental populations. For this, 500µL of saturated culture were added to 50mL of sporulation medium (1% potassium acetate, 0.05% glucose, 0.1% yeast extract, and H_2_O to 1L) and left for 5 days with gentle shaking at room temperature. We then mixed 500µL of each of the two parent spore cultures together in a 1.5mL Eppendorf tube, and centrifuged the culture down to a pellet to induce mating. To select for F1 hybrids, we inoculated 25µL of the pellet in 5mL of -lys SD medium with G418 (0.67% bacto-yeast nitrogen base without amino acids, 2% glucose, dropout powder (-lys), G418 400mgL^-1^ and H_2_O to 1L). This selects for F1 hybrid genotypes in bulk, which are heterozygous for both selective markers (-lys and G418). Note that this mass mating and sporulation protocol permitted F1 hybrid populations to contain genetic diversity, depending on how many different parental genotypes participated in the mass mating. To generate F2 hybrid populations, we put F1 hybrid populations through one round of meiosis by mass sporulating them in 50mL of sporulation medium for 5 days with gentle shaking at room temperature. F1 AND F2 hybrid populations were stored at −70°C in 22% glycerol solution for whole genome sequencing.

### Assessing parental phenotypic divergence

We phenotypically profiled all parent populations (i.e. populations evolved in NaCl, LiAc 0.01M, LiAc 0.02M, and EtOH) in 90 different environments, at five time points of divergence (at 100, 300, 500, 700, and 1000 generations), using the Scan-o-Matic system for high-throughput microbial phenotyping (Zackrisson et al. 2016; Li et al. 2019; Alalam et al. 2020; Mukherjee et al. 2021; Persson et al. 2022; Stenberg et al. 2022) version 2.2 (https://github.com/Scan-o-Matic/scanomatic.git). Parental populations were precultured in SC media for 48h in 96-well plates at 30°C without shaking and frozen. Then, cultures were pinned directly from thawed cryo-stocks in 96-format for preculturing on agar plates (for more detailed methods see (Persson et al. 2024)). Samples were split into duplicates (two 96-well plates). A third plate was used for propagation of a spatial control, containing a Y55 diploid strain. After preculture, populations were pinned onto 1536 colony arrays on agar plates robotically (Singer ROTOR HDA), allowing for accurate measuring of colony growth using the Scan-o-matic software. The spatial control was placed in every fourth position (in total 384 colonies per plate) to remove any spatial bias, irregularities in the plates, or other bias from the local area. In total, 52 distinct stressors were used in different concentrations, resulting in 90 distinct environments, including superoxidative stress (e.g. paraquat), heavy metals (e.g. CuSO_4_), different carbon sources (e.g. maltose), antifungal drugs (e.g. Clotrimazole). Phenotypic data, i.e. growth measurements, are reported in Table S1, with the full list of environments in Table S2. Cultivation plates were maintained undisturbed and without lids for the entire duration of the assay (72h) in high-resolution desktop scanners (Epson Perfection V800 PHOTO scanners, Epson Corporation, UK) at 30°C, in moisture-controlled thermostatic cabinets with air circulation. We imaged plates at 20-minute intervals using transmissive scanning at 600 dpi and extracted intensities for pixels included in, and outside, each colony. We estimated the sum pixel intensity for each colony, subtracted the median pixel intensity of the local background, and transformed the remaining cell-associated pixel intensity to cell counts by using a pre-established calibration function, established using both spectrometry and flow cytometry (Zackrisson et al. 2016). We smoothed and quality-controlled growth curves, rejecting approximately 0.3% of growth curves as erroneous, blind with respect to sample identities. Based on the number of cells after 72h, we used the yield normalized against the spatial control (Y55 diploid).

A custom ‘ParentalDivergence’ function was generated in R, using normalized yield, the number of generations in experimental evolution, and the type of population comparison. Mean and double standard deviations of normalized yields were first computed for each of the 90 environments for each of the replicate parent 1 (P1) and 2 (P2) populations. This was done for each type of population comparison, i.e. divergent-environment comparisons (NaCl x EtOH, NaCl x LiAc0.01, NaCl x LiAc0.02) and parallel-environment comparisons (NaCl x NaCl, EtOH x EtOH, LiAc0.01 x LiAc0.01, LiAc0.02 x LiAc0.02). We considered fitness to be higher if the normalized mean yield of P2 was larger than the normalized mean yield of P1 plus P1’s double standard deviation (i.e. mean P2 > mean P1 + 2σ). We considered fitness to be lower if the normalized mean yield of P2 was lower than the normalized mean yield of P1 minus P1’s double standard deviation, following previous protocols (i.e. mean P2 < mean P1 - 2σ; see (deVicente and Tanksley 1993; Brice et al. 2021). Parental phenotypic divergence is thus defined as the proportion of the 90 environments in which there was a large difference in yield between the two populations compared.

### Assessing reproductive isolation

To measure productive isolation between the parental yeast populations evolving in divergent and parallel environments, we mated together individual parent genotypes to produce F1 crosses, after 100, 300, 500, 700 and 1000 generations of experimental evolution. F1 were then sporulated to produce F1 spores (gametes), which are their recombined, haploid F2 offspring, and reproductive isolation was measured as the fraction of unviable F2 gametes. For this, F1 crosses were plated onto solid YPD agar medium (1% yeast extract, 2% peptone, 2% glucose, 2% agar made up to 1L with sterilized deionized H_2_O) and grown for 2 days, followed by replica plating onto solid KAc medium (1% potassium acetate, 0.05% glucose, 0.1% yeast extract and H_2_O to 1L) for sporulation. Populations were left for 2-7 days, until tetrads were identified under the microscope. Yeast tetrads usually contain four spores (gametes), which are the recombined haploid products of meiosis. Sporulated cultures with tetrads were incubated for one hour at room temperature in 10µL of filter sterilized zymolyase solution (1mg of zymolyase per 1mL of 1M sorbitol solution) to kill vegetative (unsporulated) cells and digest the outer membrane of the tetrads, surrounding the ascospores. Cell lysis was terminated by adding 500µL of sterilized H_2_O. To assess the fertility of F1 hybrid crosses, we dissected 10-20 individual F1 tetrads, and measured the survival rates of their haploid F2 offspring. Tetrads were dissected on YPD plates using a Singer MSM 400 robot. Individual spores were then left to germinate and grow for 72h at 30°C, at which point spore viability was assessed. Spores were counted as viable based on the presence of colonies visible to the eye. Thus, gamete viability was assessed as the capacity to germinate and show sufficient growth to form macroscopic colonies, which is a standard method in yeast genetics. The total proportion of unviable spores per cross was calculated from 20 dissected tetrads (80 spores) at 100, 700, and 1000 generations, and 10 tetrads (40 spores) at 300 and 500 generations of parental divergence.

To assess the F1 fertility of the control crosses, i.e. crosses between replicate parental populations that evolved in parallel environments, we used the same mating, sporulation and dissection protocols as described above. The total proportion of unviable spores (gametes) per cross was calculated from 10 dissected tetrads (40 spores) at 100, 500, 700 and 1000 generations of parental divergence.

### DNA extraction and whole-genome sequencing

DNA was extracted as described in (Ament-Velásquez et al. 2022). We extracted DNA from whole parental, F1, and F2 hybrid populations, respectively, at seven time points each (0, 100, 200, 300, 400, 500, 700, and 1000 generations). Extraction was carried out on overnight culture, with ∼1mL of culture used. Library preparation and sequencing were performed at the Science for Life Laboratories (Stockholm, Sweden), using the Illumina Nextera Flex protocol. Sequencing was done using either NovaSeq S4-300 (2×150bp) or NovaSeq S-Prime (2×150bp) to an estimated median depth of coverage per sample of around 30x to 780x. The raw sequences were deposited in the European Nucleotide Archive, parental populations are available at accession number PRJEB46680, while hybrids are available at accession number PRJEB83568.

### Sequencing read quality and genome alignment

FastQC v.0.11.9 (Andrews 2010) was used to assess the quality of raw sequencing reads and reads were trimmed using Trimmomatic v0.36 (Bolger et al. 2014). Sequencing reads were aligned to a S288C reference genome (R64; GenBank accession GCA_000146045.2) using BWA mem v0.7.17-r1188 (Li and Durbin 2009) and SAMtools v1.11 (Danecek et al. 2021) was used to sort and index the resulting mapping file. Duplicated reads were flagged and removed using Picard v3.1.0 (http://broadinstitute.github.io/picard/), followed by quality assessment using Qualimap v2.3 (Okonechnikov et al. 2016).

### Assessing the genomic mutational landscape

To assess the mutational landscape of each experimentally evolved, parental yeast population and the F1 and F2 crosses made between them, we measured the amount of variants detected across mutational scales, ranging from single nucleotide variants to entire chromosome aneuploidies. To identify small mutational variants (SNV and indels), we used freebayes v1.3.6 (Garrison and Marth 2012) with default parameters. Identified variants were filtered using VCFtools v0.1.16 (Danecek et al. 2011) to include only variants with minimum quality (minQ) >30 and sequencing read depth (minDP) >5. SnpEff v5.2 (Cingolani et al. 2012) was used to assess and annotate the effect of identified mutational variants on genes using the database of the reference genome R64-1-1.105 with canonical transcripts. To identify structural variants including insertions, deletions, inversions, tandem duplications and interchromosomal translocations we used Manta v1.6.0 (Chen et al. 2016) with default parameters. Structural variant detection is limited by the read pair fragment size. To more comprehensively assess the genome-wide copy number variation (amplifications and deletions), we used Control-FREEC v11.6 (Boeva et al. 2011; Boeva et al. 2012) with baseline ploidy = 2, a break-point threshold = 0.8 and coefficient of variation = 0.5 (Scopel et al. 2021). We further optimized the CNV detection by limiting the expected GC content to between 0.35 and 0.55. Variants were further filtered to only include copy number calls that were statistically significant (p<0.05) using Wilcoxon Rank Sum Test and a Kolmogorov Smirnov Test. Two metrics of CNVs were assessed for differences across time and between populations: CNV load and the fraction of the genome altered. CNV mutational load is the number of CNV events (amplification or deletion) detected within a population. The fraction of genome altered incorporates the size of each CNV event to determine the amount of the genome that is altered by CNVs. We also differentiated between small (<10kb) and large (>10kb) CNVs. Chromosomes were further considered aneuploid if >60% of the chromosome was identified as amplified or lost. To better assess the differences that might have existed between the two founding populations and potentially their lasting effects through time, we re-analyzed sequencing data from the original study (Ament-Velasquez et al. 2022), which was focused on the genetic basis of evolutionary adaptation of these founders. This study included sequencing data collected at shorter time-points, in particular within the first 100 generations (30 and 60 generations). We analyzed the relative sequencing read depth to determine how persistent the aneuploidies identified in the founding populations (generation 0) were during adaptation.

### Calculating population differentiation (F_st_) and genome similarity (Jaccard index)

We calculated pairwise F_st_ between all divergent- and parallel-evolving parental populations at each of the sampled timepoints, based on single nucleotide polymorphisms (SNPs). We first combined the vcf files generated from freebayes using bcftools v1.19 (Danecek et al. 2011), with parallelization of file compression by gnuparallel v20230422 (Tange 2011), using the default parameters of bcftools merge and the R package poolfstats v2.2.0 (Gautier et al. 2022) on R v3.4 (R Core Team 2023). First the vcf files were converted to pooldata objects, using vcf2pooldata with the default parameters. F_st_ was then calculated using pairwiseFST with the nsnp.per.bjack.block = 150 to calculate the Block-Jackknife estimation of standard error, using an ANOVA framework (Hivert et al. 2018). Similarities between copy number variation calls in different populations and through time were assessed based on the Jaccard similarity coefficient using CNVmetrics R package v3.20 (Belleau et al. 2021; Deschênes et al. 2022).

## Results

We experimentally evolved yeast populations for 1000 generations in divergent and parallel environments. The four selection environments contained either salt, ethanol, or two different concentrations of lithium acetate, and significantly differed in the degree to which they reduced the fitness of the founder populations. We have previously provided phenotypic and genetic evidence that these populations become differentially adapted to these environments (Ament-Velásquez et al. 2022). At five time points during experimental evolution, we made crosses between the parallel- and divergent-evolving populations (the ‘parents’) to measure postzygotic reproductive isolation (F1 spore viability). At seven time points, we used whole population genome sequencing on parents, F1, and F2 hybrids, to assess their mutational load, i.e. the number and types of structural variation, and other genomic features.

### Parental phenotypic divergence

We phenotypically profiled the evolving parent populations in 90 non-selection environments. These environments contained the same medium (SC) as the selective environments used for experimental evolution, but they also each contained an additional growth limiting substance (Table S2). Thus, our measure of phenotypic divergence between populations is likely a conservative estimate. Note that we only assessed the phenotypic divergence of evolving parent populations here (not the F1 or F2 crosses), to test for adaptation as a result of divergent ecological selection in the parents.

As expected, with increasing generations of experiment evolution in different environments, divergent-selected parent populations significantly diverged in phenotype (regression across all divergent-evolved populations: R = 0.14, p = 0.007; when removing the outliers at G1000: R = 0.09, p = 0.036; **Figure 2A**). Regression of individual environment comparisons resulted in significantly increasing phenotypic divergence in the NaCl vs. EtOH-evolved populations (R = 0.41, p=0.004), but not for the other two comparisons.

**Figure 2:**
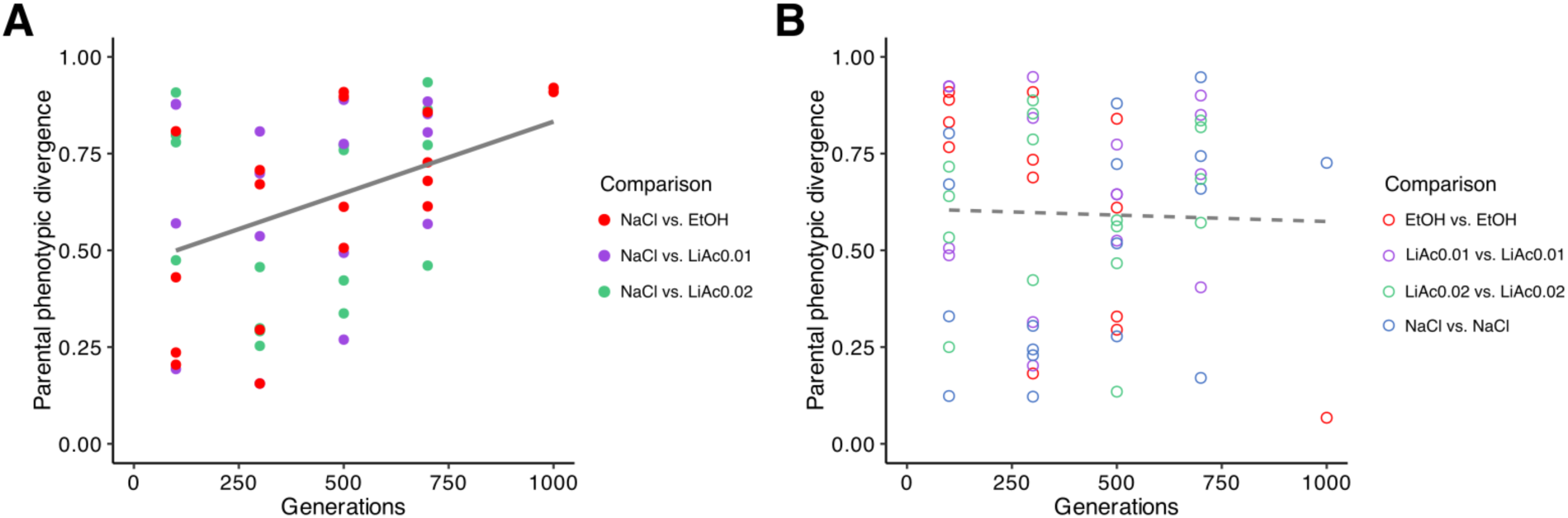
Phenotypic divergence of evolving parental populations. Parental phenotypic divergence (the proportion of 90 environments, in which populations significantly differed in yield) at five time points (number of generations) of experimental evolution. **A)** Comparisons between divergent-evolved populations in different environments. **B)** Comparisons between parallel-evolved populations in the same environment. Solid line indicates significant regression, dashed line shows non-significant regression across all comparisons.

In contrast, parent populations evolving in parallel in the same environment did not diverge phenotypically (R = 9.34e-04, p = 0.81; when removing the outliers at G1000: R = 0.002, p = 0.768; **Figure 2B**). Here, individual regression of the EtOH vs. EtOH-evolved populations resulted in a significant decrease of phenotypic divergence with generations (R = 0.51, p = 0.006) but this trend was no longer significant when removing the outlier at G1000 (R = 0.30, p = 0.066). This suggests that the phenotypic divergence of populations evolving under different selection pressures was caused by ecological differences between the four selection environments.

### Genetic population differentiation

When comparing parental populations, genetic population differentiation (genome-wide F_st_) was generally larger (mean = 0.25) in pairwise comparisons between divergent-selected (grey boxes in **Figure 3A**) than between parallel-selected populations (mean = 0.19, colored boxes). The only exception were comparisons between parallel-selected LiAc0.01 populations (purple), which produced slightly higher F_st_-values but were not significantly larger than the divergent-selected populations. F_st_ -values increased with the number of generations of evolution in both divergent and parallel environments (**Figure 3A**, Figure S1), except for comparisons between divergent-selected NaCl and LiAc0.02 populations. NaCl and LiAc0.02 populations tended to decrease in F_st_ over time. This may be the result of populations losing genetic variation overall, which was particularly the case for LiAc0.02 populations as discussed below. Importantly, the two founding populations with compatible selective markers were found to have a very low pairwise F_st_-value suggesting a high level of genetic similarity and low amount of differentiation (Figure S2).

**Figure 3:**
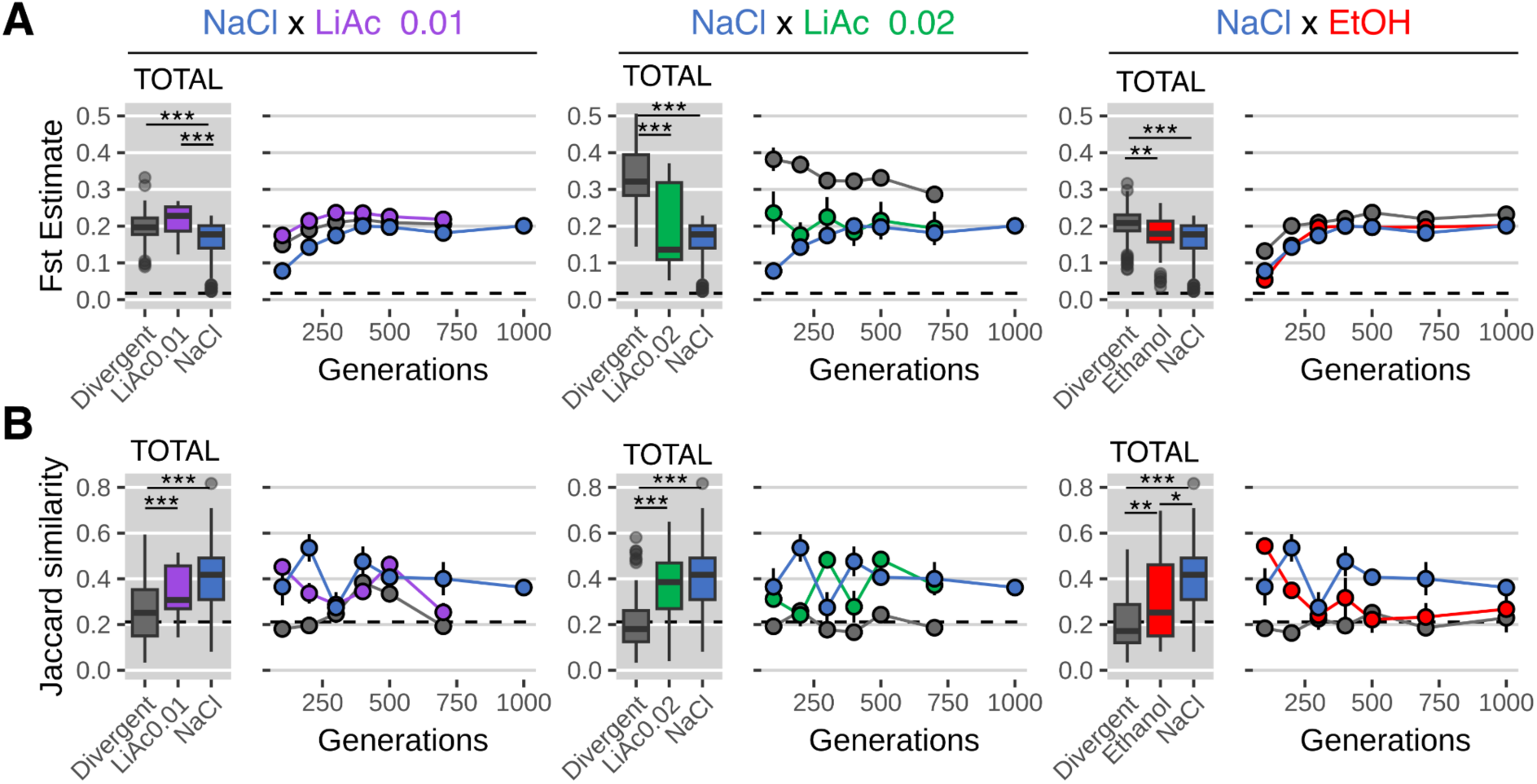
Parental genomic population differentiation. **A)** Genetic population divergence (F_st_) at seven time points (n of generations) of experimental evolution. . Boxplots indicate the first and third quartiles with whiskers extending to the furthest point not exceeding 1.5 x the interquartile range and outliers beyond this are shown as dots. The solid line indicates the median value. **B)** Genomic similarity (Jaccard similarity index) at seven time points (n of generations) of experimental evolution. Box plots show means (thick horizontal line) across all generations of each population type. Circles indicate the mean measurement and the vertical line indicates the standard error of the mean. ‘Divergent’ indicates comparisons between divergent-selected populations. Asterisks indicate statistical significance using Kruskal-Wallis and pairwise Wilcoxon post hoc tests with Bonferroni corrections (adjusted p-values < 0.001 = ***, < 0.01 = **, < 0.05 = *). Dashed horizontal lines indicate the measured differentiation between the founder populations at the beginning of evolution.

Consistent with their lower F_st_ -values overall, the genomes of parallel-selected populations remained significantly more similar (using the Jaccard similarity index) than the genomes of divergent-selected populations (gray boxes in **Figure 3B**). Together, this suggests that the larger genetic divergence of populations evolving under different selection pressures was caused by ecological differences between the four selection environments.

### F2 hybrid viability

The viability of F1 spores made between divergent-selected populations was significantly lower than the viability of crosses made between parallel-selected populations on average, when analysed across all cross types and all time points (Wilcoxon rank sum test with continuity correction; W = 1395.5, p = 0.046; **Figure 4**, top left hand boxplot). Analyzing cross types separately, the viability of offspring from parallel-selected populations was significantly higher than the viability of hybrids made from divergent-selected parents in two of the three cross types, when averaged across all time points (Kruskal-Wallis: NaCl x EtOH: X^2^ = 7.6216, p = 0.022; NaCl x LiAc0.02: X^2^ = 14.209, p = 0.001). In the NaCl x LiAc0.01 cross type, the viability between parallel-selected populations was similar to that between divergent-selected populations (X^2^ = 5.834, p = 0.054).

**Figure 4:**
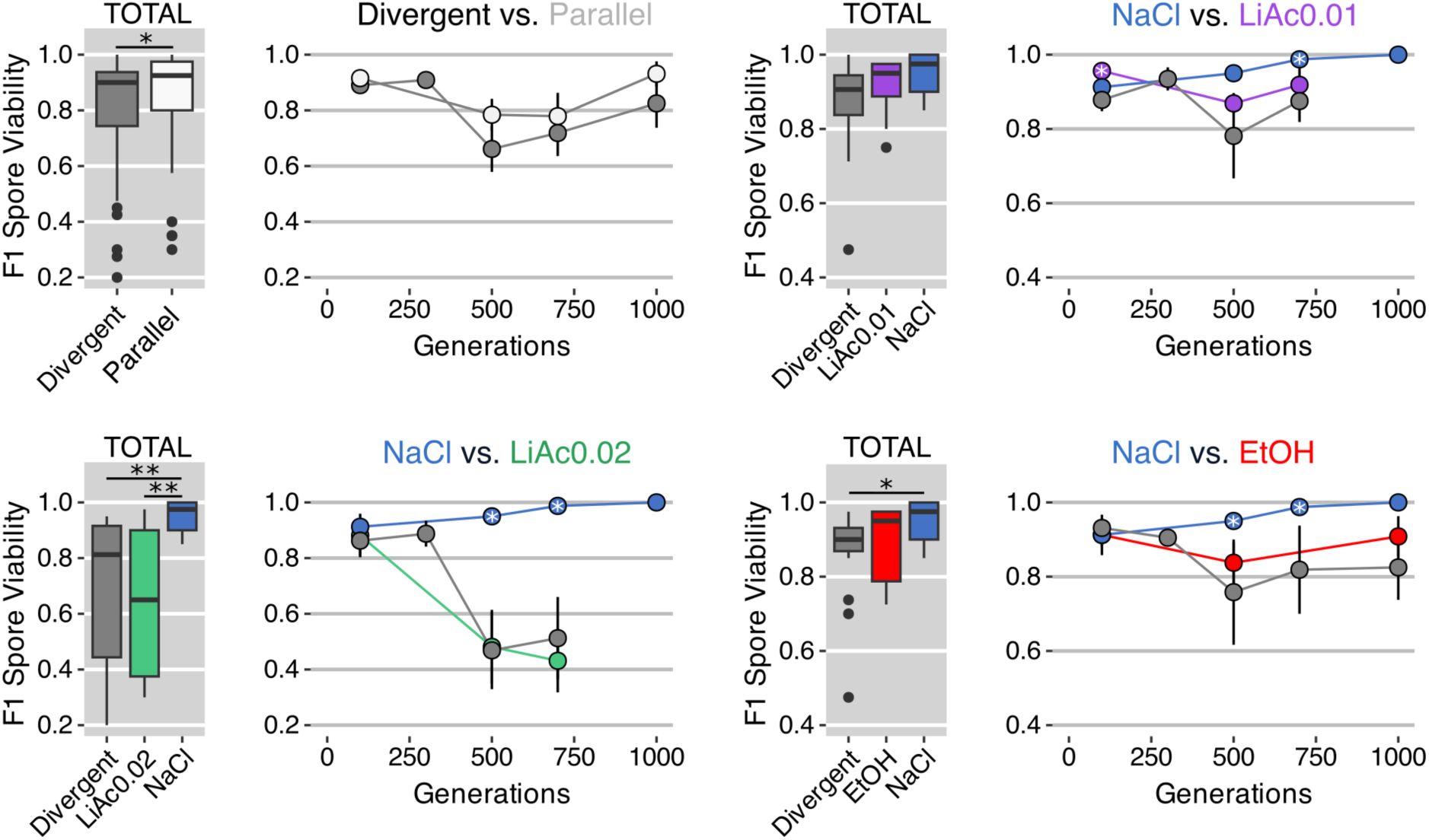
Analysis of reproductive isolation. The viability of F1 spores (i.e. the recombined haploid F2 offspring), which is an inverse measure of reproductive isolation, at five time points (number of generations) of experimental evolution. Circles indicate the mean measurement and the vertical line indicates the standard error of the mean. Crosses made between parallel-selected populations evolving in the same environment are coloured (or white in the top left hand plot), crosses made between divergent-selected populations, evolving in different environments, are grey. Box plots show means (thick horizontal line) across all generations of each cross type. ‘Divergent’ are the crosses between divergent-selected populations (NaCl vs LiAc0.01, NaCl vs LiAc0.02, NaCl vs EtOH). Black asterisks indicate statistical significance using Kruskal-Wallis and pairwise Wilcoxon post hoc tests with Bonferroni correction (adjusted p < 0.001 = ***, < 0.01 = **, < 0.05 = *). White asterisks inside circles indicate significantly lower viability in divergent-than in parallel-selected crosses, using one-sided Wilcoxon tests with Bonferroni corrections (adjusted p < 0.05 = *).

Following the breakdown of hybrid viability over time, the viability between divergent-selected parents was still relatively high at early time points of divergence (on average 89.1% ± 0.02 at 100 generations), but then decreased substantially between 250 and 500 generations to on average 66% ± 0.08 (**Figure 4**, top left hand line plot) below that of parallel-selected crosses. This pattern was not consistent among all cross types. The viability of two parallel-selected cross types (LiAc0.01 x LiAc0.01, EtOH x EtOH) was higher than that of the divergent-selected crosses on average (although not all within-time point comparisons were significant), but a notable exception were the crosses made between the parallel-selected LiAc0.02 x LiAc0.02 populations, which dropped in viability after 500 generations, to a similar extent as that of the divergent-selected NaCl x LiAc0.02 hybrids. We discuss potential explanations for this below.

### The genome-wide mutational landscape of adaptation and hybridization

In an effort to find the genetic mechanisms potentially underlying postzygotic reproductive isolation, we assessed the genome-wide mutational landscape of the divergent-evolved parental populations, and the F1 and F2 hybrid crosses made between them, at all seven sampled timepoints. We focused on three major mutational classes: single nucleotide variants (SNVs), large structural variants (SVs), and copy number variants (CNVs).

As might be expected from selective sweeps during directional selection, we found that all populations showed reduced SNV mutational diversity compared to the founder population within the first 100 generations of evolution (dashed black lines in Figure S3A). Most identified mutations were single nucleotide variants, fewer were insertions and deletions (indels) (Figure S4). The mutational SNV load in LiAc0.02 populations was consistently lower than in populations evolving in NaCl and in their F1 and F2 hybrids, and remained low through time (Kruskal-Wallis: X^2^ = 44.95, p <0.001, Figure S3A). This pattern remained after filtering for mutational variants estimated by SnpEff to have high impact (Kruskal-Wallis: X^2^ = 13.07, p=0.004, **Figure 5A**). LiAc0.02 populations also showed a lower SNV mutational load than the populations evolving in the weaker concentration of lithium acetate (LiAc0.01). It is possible that this is the result of stronger purifying selection in LiAc0.02 populations from the start. But since populations evolving in the stronger concentration of lithium acetate have likely also undergone fewer mitotic generations, this may also reflect mutations accumulating primarily as a function of reproductive time. In the other three parental population types (NaCl, LiAc0.01 and EtOH), SNV mutational load decreased over time (**Figure 5B**) and the pattern stayed consistent when filtering for only mutations with high impact on fitness (Figure S5A). Conversely, in the F1 (R = 0.48, p < 0.01, in grey) and F2 hybrid populations (R = 0.57, p < 0.01, in yellow), we found a total increase in mutational load over time (**Figure 5B**). Thus, the more the parental populations diverged phenotypically and genetically over time, the more single nucleotide mutational variants were detected in their resulting F1 and F2 hybrid populations.

**Figure 5:**
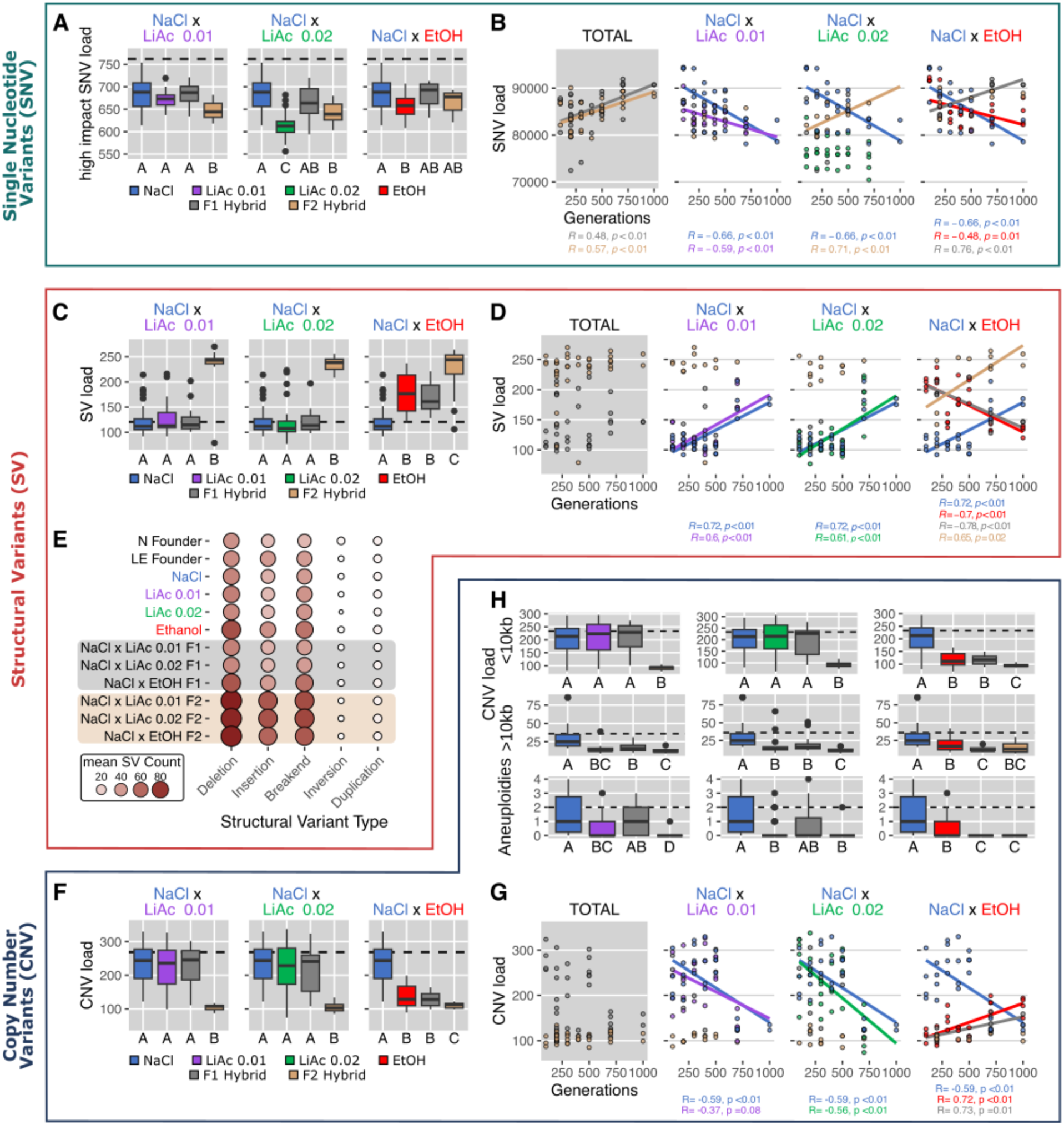
The genome-wide mutational landscape of hybridization at seven time points of experimental evolution. **A)** Single nucleotide variant (SNV) mutational load after filtering for high impact mutations on gene function using SnpEff v5.2 (Cingolani et al. 2012). The mutational load is averaged across all timepoints for each hybrid cross. The mean mutational load of the founder population is indicated by the dashed black line. Boxplots indicate the first and third quartiles with whiskers extending to the furthest point not exceeding 1.5 x the interquartile range and outliers beyond this are shown as dots. The solid line indicates the median value. Different letters (A, B and C) indicate statistically significant differences using Kruskal-Wallis and pairwise Wilcoxon post hoc tests with Bonferroni corrections. **B)** Linear regression (Pearson R) of SNV mutational load at seven time points (n of generations) of experimental evolution. Only significant linear relationships are shown (p ≤ 0.05). The gray plot includes F1 and F2 hybrids across all crosses. **C)** Structural variant (SV) mutational load averaged across all sampled timepoints (TOTAL) for the three types of hybrid crosses. SV mutational load consists of the number of deletions, insertions, inversions, tandem duplications and interchromosomal translocations (> 8bp) detected in the population. Colors, boxplots, and statistical significance are consistent with panel A. **D)** Linear regression (Pearson R) between SV mutational load and increasing timepoints of parental divergence (number of generations). Only significant linear relationships are shown (p ≤ 0.05). The gray plot includes F1 and F2 hybrids from all crosses. **E)** Mean structural variant count of each type of structural variant (deletion, insertion, breakend, inversion and duplication) for each environment and cross. Node color and size indicates the mean count of each variant type across all timepoints. **F)** Copy number variant (CNV) mutational load consisting of amplifications, deletions and aneuploidies. CNV mutational load averaged across all sampled timepoints (TOTAL) for the three types of hybrid crosses. Colors, boxplots, and statistical significance are consistent with panel A. **G)** Linear regression (Pearson R) between CNV mutational load and increasing timepoints of parental divergence (number of generations). Only significant linear relationships are shown (p ≤ 0.05). The gray plot includes F1 and F2 hybrids from all crosses. **H)** The distribution of large (>10kb) CNVs, small (< 10kb) CNVs and chromosome aneuploidy (> 60% amplified or deleted) across all sampled timepoints for the three types of hybrid crosses. Colors, boxplots, and statistical significance are consistent with panel A.

Analysis of structural variants (i.e. insertions, deletions, duplications, inversions and interchromosomal translocations that are > 8bp) showed that the total number of SVs found in the evolving parental populations remained comparable to the levels in the founder population (dashed black lines in **Figure 5C**). Unlike the pattern we found in SNV mutational load, the number of SVs harbored in parental populations increased over time (**Figure 5D**). This positive linear relationship was consistent for populations evolving in NaCl (R = 0.72, p < 0.01), LiAc0.01 (R = 0.6, p < 0.01), and LiAc0.02 (R = 0.61, p < 0.01), but not for EtOH populations, which already had elevated levels of SVs (p < 0.001) early on but then decreased over time (R = −0.7, p < 0.01; Figure S6A).

Strikingly, we found very high levels of SVs in all types of F2 hybrid crosses, right from the beginning of parental population divergence (Kruskal-Wallis: NaCl x LiAc0.01: X^2^ = 22.87, p <0.001; NaCl x LiAc0.02: X^2^ = 30.28, p <0.001; NaCl x EtOH: X^2^ = 34.45, p <0.001). In most cases, F2 hybrid populations harboured almost twice as many structural variants compared to the corresponding parental populations and the F1 hybrids. This pattern was seen across SV types collectively, but was particularly prominent for insertions, deletions and interchromosomal translocations (breakend mutations, **Figure 5E**, Figure S7-S11). Duplications were rare compared to deletions and insertions and are therefore not driving the elevated levels of SVs observed in F2 hybrid genomes (**Figure 5E**). Because F2 hybrids are distinguished from F1 hybrids and the parent populations only by having passed through one round of meiosis, the additional load of structural variants we found in F2 hybrids likely arose due to meiotic errors.

In addition to structural variants, we also more comprehensively assessed copy number variations (CNV) across the genome, which are a subtype of structural variation. In the structural variation analysis, detection of variants was limited by the read pair fragment size. However, copy number variants can range from small focal amplifications or deletions to chromosome arm, or whole chromosome aneuploidies. Therefore, we used an approach that employs sequencing read depth and allelic imbalances to estimate segmented copy number uniformly throughout the genome. On average, parental populations, F1 and F2 hybrids had a lower CNV mutational load compared to the founder population (dashed black lines in **Figure 5F**), suggesting that directional adaptation has reduced CNV load in all environments. This pattern remained consistent when including the size of each copy number variant to estimate the fraction of altered genome content (Figure S12B). Most populations showed moderate levels (< 10%) of total alteration (amplification + deletion) of the genome by CNV (Figure S12B). However, NaCl populations had a larger fraction of their genomes altered in copy number than parental populations evolved in LiAc and EtOH (pairwise Wilcoxon test: vs LiAc0.01: p <0.001; vs LiAc0.02: p <0.001; vs EtOH: p =0.012). This is due to three parallel-evolved NaCl populations that had a large fraction of genome alterations (> 20%) at three different time points of evolution.

Strikingly, the F2 hybrid genomes of all three crosses contained significantly fewer CNVs than parental populations and most F1 hybrids (**Figure 5F**; Kruskal-Wallis: NaCl x LiAc0.01: X^2^ = 30.1, p < 0.001; NaCl x LiAc0.02: X^2^ = 24.4, p < 0.001; NaCl x EtOH: X^2^ = 37.7, p < 0.001) and F2 hybrid genomes of all crosses were significantly less altered by CNVs (in % genome) than the parental populations (Figure S12B). Overall, we found that amplifications were more common than deletions across populations and through time (Figure S13).

Similar to SNVs and SVs, we found consistent patterns of increase or decrease in CNVs with the number of parental generations of evolution in divergent environments (**Figure 5G**). In the parental NaCl (R = −0.59, p < 0.01) and LiAc0.02 (R = −0.56, p < 0.01) populations, the CNV mutational load decreased with the number of generations. In the LiAc0.01 parents, the same pattern was seen but was not significant (R = −0.37, p = 0.08). Interestingly, CNV mutational load was often relatively stable until approximately 500 generations, followed by a steep decrease (**Figure 5G**, Figure S12A). In the EtOH parents, we found the opposite pattern, with CNV load increasing with the number of generations (R=0.72, p <0.01), affecting the F1 hybrid genomes between NaCl and EtOH in the same way (R = 0.73, p = 0.01).

Lastly, we sought to understand if these patterns of CNV load were driven by small focal variants or larger multi-kilobase alterations, including whole chromosomal aneuploidies. We found that the majority of variants were small (< 10kb) and contributed significantly to the global pattern found in CNV load (compare upper row of plots in **Figure 5H** to **Figure 5F**). A closer examination of the number of larger alterations (> 10kb; middle row of plots in **Figure 5H**), revealed that the F2 hybrid genomes of two hybrid crosses (NaCl x LiAc0.01, NaCl x LiAc0.02), showed significantly fewer large alterations (on average 12.1) than their respective parents (on average 20.8) and F1 hybrids (on average 19.4). This pattern is further emphasized when examining exclusively the number of aneuploid chromosomes (>60% amplified or deleted) found in each sample (lower row of plots in **Figure 5H**). On average, F2 hybrid genomes carried fewer aneuploidies (mean=0.24 chromosomes) compared to parent (mean=0.67) and F1 hybrid genomes (mean=0.64).

### The distribution of mutational events across the genome

After assessing the genome-wide mutational landscape in terms of SNVs, SVs, and CNV mutational load, we determined the distribution of these mutational events across the genome. We found that the rate of SNVs was not uniformly distributed, but instead showed chromosomal biases (Kruskal-Wallis: X^2^ = 147.28, p <0.001, **Figure 6A**). Chromosomes VI and VIII had significantly higher frequencies of SNVs (16.7 and 14.1 mutations/kb, respectively). Chromosome I and IX also showed slightly elevated SNV frequencies (10.8 and 9.35 mutations/kb, respectively). Chromosome VII on the other hand showed a significantly lower frequency of SNVs (2.29 mutations/kb). The remaining chromosomes had approximately equal frequencies of SNVs (6.39-7.34 mutations/kb). These chromosomal biases in SNV distribution already existed in the founder population and remained consistent across all parental lineages and all F1 and F2 hybrids (**Figure 6A**).

**Figure 6:**
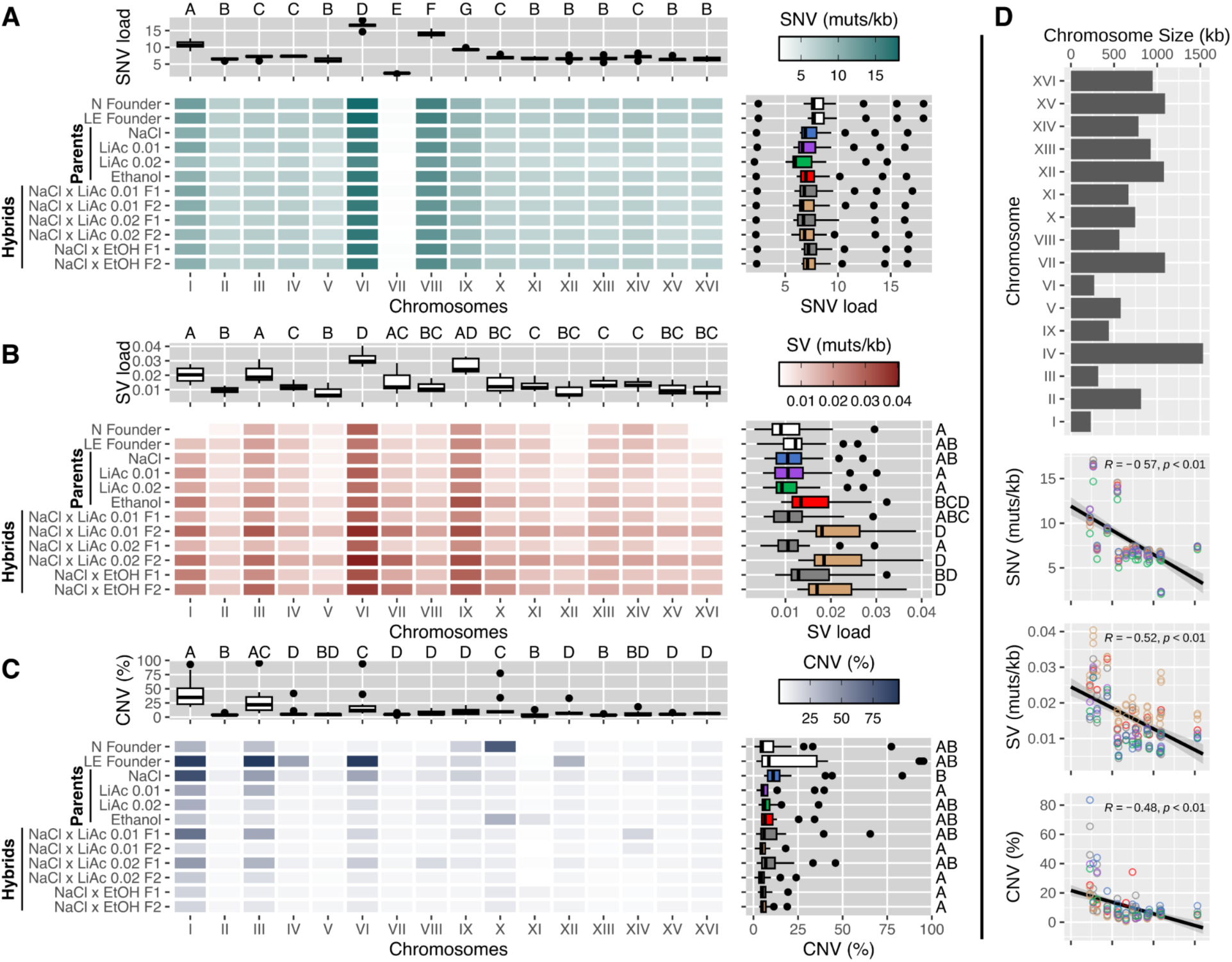
The distribution of mutational events across the genome. **A)** The mean number of events (mutations/kb) for each chromosome across all available timepoints. The top graph shows the distribution of mean SNV counts for each chromosome, while the right graph shows the distribution for each environment and hybrid cross. Boxplots indicate the first and third quartiles with whiskers extending to the furthest point not exceeding 1.5 x the interquartile range and outliers beyond this are shown as dots. The solid line indicates the median value. Different letters (A, B, C, D) indicate statistically significant differences using Kruskal-Wallis and pairwise Wilcoxon post hoc tests with Bonferroni corrections. **B)** The mean number of structural variant (SV) events (mutations/kb). **C)** The mean amount of genome altered (%) by copy number variants (losses and gains), including whole chromosome aneuploidies. **D)** The relationship between chromosome size and SNVs, SVs or CNVs. The barplot indicates the size of each of the 16 chromosomes in *Saccharomyces cerevisiae* reported in kilobases. The linear correlations show the mean SNV (muts/kb), SV (muts/kb) or CNV (% genome altered) for each population and for each chromosome as shown in the top graphs of panels A, B and C. The colors of each node indicate the populations (NaCl, LiAc0.01, LiAc0.02, EtOH, Hybrid F1, Hybrid F2).

The distribution of structural variant (SV) events were also not uniformly distributed throughout the genome (Kruskal-Wallis: X^2^ = 60.06, p <0.001, **Figure 6B**). Similar to the SNV distribution, we found highly elevated rates of SVs on chromosome VI (0.03 mutations/kb). We also found peaks in the rate of SVs in chromosomes I, III, and IX in most parental and hybrid populations.

The proportion of whole chromosomes altered by copy number variants was not consistent across environments and hybrid crosses (**Figure 6C**). Interestingly, F2 hybrids of all crosses showed the lowest levels of copy number alterations, and aneuploidy (defined as > 60% of chromosome amplification) was consistently rare across all F2 hybrid chromosomes. We discuss the implications of the near absence of aneuploidies and large genome alterations in F2 hybrid genomes below. Our analysis shows that the two founding populations each carried aneuploid chromosomes. The NaCl founder population (N founder, **Figure 6C**) had an extra chromosome X, while the LE founder population had three aneuploid chromosomes (I, III and VI). It is important to note that deeper analysis of sequencing read depth of early generational timepoints from an earlier study (Ament-Velásquez et al. 2022) revealed that these aneuploidies were lost in the population within the first 30 or 60 generations of adaptation (Figure S14). Parental NaCl populations subsequently lost the extra chromosome X over the course of experimental evolution, but evolved an aneuploid chromosome I and underwent alterations of large portions of chromosome III and VI. Chromosome I was also heavily altered in parent populations evolving in LiAc 0.01 (40%), LiAc 0.02 (37%), and EtOH (25%). Similarly, F1 hybrid crosses between NaCl and the two LiAc populations (to a lesser extent the NaCl x EtOH F1 hybrids) also showed large copy number alterations in chromosomes I, III, and VI (**Figure 6C**). Confirming our previous result (Ament-Velásquez et al. 2022), we also recovered a chromosome X aneuploidy in two parental EtOH populations and the chromosome XI aneuploidy in one parental EtOH population here.

The 16 chromosomes of *S. cerevisiae* range widely in size from 270kb (chrVI) to 1,531kb (chrIV; **Figure 6D**). As might be expected, larger chromosomes are larger targets for SNVs, SVs and CNVs and therefore have higher overall mutational load for each mutational class. However, after correcting mutational load for chromosome size, either by reporting SNVs and SVs as mutations per kilobase or reporting CNVs as the percentage of chromosome altered, we found a strong negative relationship between chromosome size and SNV, SV and CNV mutational rate (**Figure 6D**). Smaller chromosomes had elevated rates of mutation while larger chromosomes were rarely affected. This pattern is most striking in CNVs where, as previously mentioned, chromosomes I, III and VI were disproportionately altered. This is interesting because chromosomes I, III, and VI are the smallest chromosomes in the yeast genome and have been frequently reported to carry aneuploidies in other yeast studies as well (reviewed in (Gilchrist and Stelkens 2019), likely because aneuploidies of smaller chromosomes cause fewer deleterious gene dosage effects and can be compensated for at the proteome level (Muenzner et al. 2024). This pattern of chromosome size bias for CNVs extends beyond intraspecific crosses and has been found across interspecific yeast hybrids with diverse ancestries (Bendixsen, Peris, et al. 2021).

## Discussion

We used experimental evolution with budding yeast to test for the evolution of reproductive isolation as a result of ecologically divergent selection, a process called ecological speciation (Mayr 1947; Schluter 2001; McKinnon et al. 2004; Nosil 2012; Jarrett et al. 2025). Previous studies with yeast have tested for reproductive isolation between species that have diverged millions of years ago, which translates into billions of generations (Greig et al. 2002; Shapira et al. 2014; Stelkens et al. 2014; Bernardes et al. 2017; Eberlein et al. 2019; Gallone et al. 2019; Zhang et al. 2020; Bautista et al. 2021; Brice et al. 2021; Bautista et al. 2023). Here, instead of starting with already divergent species, separated by billions of generations, each with a complex evolutionary history of selection, genetic drift, recombination, and gene flow, we generated phenotypic and genetic divergence over only a few hundred generations. We evolved replicate populations in allopatry, in four different environments, under both ecologically divergent and parallel selection. Hence, we controlled the type and degree of selection, minimized genetic drift and recombination, and excluded gene flow. We systematically produced hybrid crosses between these populations at increasing time points of experimental evolution, to test whether reproductive isolation evolved between the divergent-selected populations, as predicted by ecological speciation theory.

### Divergent ecological selection leads to partial reproductive isolation

First, we confirmed that populations when evolved in different environments, increasingly diverged in phenotypic traits that were not directly under selection. The parallel-selected (control) populations on the other hand remained phenotypically similar (**Figure 2**). This provides evidence for large scale phenotypic differentiation under divergent selection. Second, the genomes of divergent-selected populations became more genetically differentiated over time, showing higher F_st_ -values and lower similarity (Jaccard index) than those of the parallel-selected populations (**Figure 3**). Third, we found that divergent-selected populations produced hybrid gametes with reduced viability, which is a classic signature of postzygotic reproductive isolation in the form of hybrid breakdown (**Figure 4**). Zygotes generated from parallel-selected populations on the other hand remained reproductively more compatible, with some exceptions (**Figure 4**). Together, our results suggest that partial reproductive isolation evolved due to divergent ecological selection in our experiment. Alternative processes, such as mutation-order effects (Schluter 2009; Ono et al. 2017), may have contributed to the build up of reproductive barriers, but since most parallel-selected populations remained compatible, neither the timing nor the order in which mutations occurred likely played a large role.

### Recombination errors may cause genomic instability in hybrids

Early experiments, e.g. in fruit flies with similar set-ups of divergent-and parallel-selected lines on different media (Dodd 1989) and a large body of theory (reviewed in Kirkpatrick and Ravigné 2002) have demonstrated the evolution of reproductive isolation as a consequence of adaptive divergence (Nosil 2012). Here, we have taken additional steps to understand the genetic basis of this hybrid breakdown. Our population genomic analyses show that, as parental populations diverged phenotypically and genetically over time, the mutational load of single nucleotide variants (SNVs) steadily increased in F1 and F2 hybrids (**Figure 5B)**, with a bias for chromosomes I, VI, and VIII (**Figure 6A**). We speculate that the sudden breakdown of hybrid viability between 250 and 500 generations (**Figure 4**) may be a threshold effect of mutational load, where mutations have only small, non-linear effects on viability until a critical point is reached, after which DNA repair pathways are increasingly defective in hybrid genomes.

Even more strikingly, F2 hybrid populations harboured more than twice as many structural variants (SVs) than their respective parental populations and the F1 hybrids. This pattern was seen across all SV types (**Figure 5C**), but was particularly strong for insertions, deletions, and interchromosomal translocations (breakend mutations, Figure S7-S11) with strong chromosomal biases for chromosomes I, III, VI, and IX (**Figure 6B**). Because F2 hybrids differ from F1 hybrids only by one meiotic event (and a few mitotic generations before and after sporulation that are negligible), the increase in structural variants seen in F2 hybrids can only reasonably be explained by errors during meiosis. This suggests that ecological divergence has led to hybrid offspring with higher genomic instability due to meiotic recombination errors (e.g. unequal or incorrect crossing overs), resulting in frequent chromosomal rearrangements (SVs). However, additional sequencing data from crosses between parallel-selected populations (not available to us yet) would be needed for comparison, to conclusively test for causality between divergent selection and genome instability.

Interestingly, F2 hybrids showed the lowest levels of copy number alterations, and aneuploidy rates were consistently low across all chromosomes in F2 hybrid genomes (**Figure 5F**, **Figure 6C**). Meanwhile, all parent and F1 hybrid populations contained various aneuploidies, especially in chromosomes I, III, and VI, which are the smallest chromosomes in the yeast genome and frequently reported to carry aneuploidies (reviewed in Gilchrist and Stelkens 2019). Note, that the aneuploidies originally present in the founder populations (Figure S14) had disappeared from the evolving parental populations before we made the first hybrid cross at 100 generations, hence all aneuploidies arose *de novo* over the course of experimental evolution, or after meiosis in the F2 genomes. The absence of aneuploidies in F2 hybrids is surprising and contrasts with the high SV load found in hybrid genomes. We expected errors during meiosis to also lead to increased chromosome non-disjunction and aneuploidy. A plausible explanation is that, by default, we only sequenced viable genotypes that had successfully passed through meiosis and germinated. Average mortality due to lethal aneuploidies is likely high in the recombined haploid gametes, where copy number imbalances imposed by aneuploidies are more pronounced, and lethality has additional opportunities to manifest when strains are pushed through germination. As a result, aneuploidies may have been efficiently purged from F2 hybrid genomes before the sequencing stage.

### A putative genetic mechanism for reproductive isolation

Experimental and genetic work in interspecific yeast crosses, made from species separated by billions of generations and high nucleotide divergence (>10%), has shown that the main reason for hybrid breakdown in yeast is antirecombination (Hunter et al. 1996; Rogers et al. 2018; Bozdag et al. 2021). Antirecombination suppresses crossovers during meiosis when nucleotide divergence along chromosomes is too high. The ensuing mispairing of homologous chromosomes causes missegregation into the hybrid gametes, leading to lethal aneuploidies and typically > 90% spore mortality in interspecific yeast crosses. This process has also been shown to cause hybrid sterility in mice, where it is controlled by the Prdm9 gene (Forejt and Jansa 2023). However, antirecombination due to genome-wide nucleotide divergence is unlikely to have played a role in the hybrid breakdown we observed here, given the short evolutionary time span of our experiment. Instead, we suggest that the following process underlies the build-up of reproductive barriers here: First, divergent ecological selection in the parent populations has led to SVs with positive (adaptive) fitness effects and additional neutral SVs hitchhiking along with them. These parental SVs then interfered with hybrid meiosis, causing additional SVs to emerge in the hybrid genomes, due to unequal or incorrect crossing over. Cumulatively, the high SV load of the hybrid genomes then negatively affected gamete viability, germination, and colony formation, disrupting cell function and homeostasis (Chang et al. 2013; Liu et al. 2022). In short, we suggest that the most likely genetic mechanism underlying hybrid breakdown in our experiment is the high structural variant load of the hybrid genomes, which was jump-started by the increasing number of SVs accumulating in the divergent-selected parent populations (**Figure 5D**). Whether the accumulation of SVs in parent populations under divergent selection may be a common mechanism leading to genomic instability in hybrids needs to be tested in other species and systems.

We found an interesting exception in crosses between parallel-selected LiAc0.02 populations where, against our expectations, gamete viability decreased over time (**Figure 4**). We previously found that three of four replicate populations evolving in LiAc0.02 had turned (functionally) haploid early on in experimental evolution (Ament-Velásquez et al. 2022). In haploid populations, beneficial mutations can sweep to fixation faster and deleterious mutations are more quickly purged by purifying selection, as they are not masked by heterozygosity (Gerstein et al. 2011; Cervantes et al. 2023). This is consistent with our finding that the mutational SNV load in LiAc0.02 populations was consistently lower than in other populations, which may be the result of stronger purifying selection in this environment. These processes are particularly efficient in large populations exposed to stressful environments with strong selection, which the LiAc0.02 environment represents (Ament-Velásquez et al. 2022). Propelled by clonal interference, this may have affected the adaptive dynamics (the rate and direction of allele frequency changes) in the LiAc0.02 populations in idiosyncratic ways, leading to substantial genetic differentiation and incompatibilities between them, despite parallel selection pressures.

Although we may speculate on the mechanisms causing reproductive isolation in our experiment, the underlying genetic architecture of reproductive isolation remains unidentified. The pattern of decreasing genomic similarity and increasing F_st_ between divergent-selected populations (while parallel-selected populations remained similar), and the vast genome-wide instability found in hybrid genomes, does suggest that reproductive isolation is based on (or at least accompanied by) multiple genomic regions and that ‘continents’ rather than ‘islands’ of differentiated loci are operating together (Michel et al. 2010). However, due to the absence of sexual recombination during the evolution of the parent populations in our experiment, other evolutionary mechanisms may have also caused large genomic regions to become differentiated. Effects of clonal interference and linked selection for example, i.e. the competition between multiple beneficial mutations arising independently in the same population leading to only one lineage sweeping through the population, may have caused genetic differentiation between populations that is not due to ecological selection alone. Although our previous work has identified the number and types of adaptive mutations segregating and fixing in the evolving parental populations (Ament-Velásquez et al. 2022), our pool-seq approach does not provide haplotype information. Thus, we cannot infer which and how many genes are causing reproductive isolation, how large their effect sizes are, and whether they are genomically clustered (Nosil and Feder 2012). Directed crosses between parents with known genotypes, and dissecting and genotyping a large number of recombined F2 gametes, could identify incompatible chromosomes, chromosomal regions, or individual genes causing reproductive isolation in this system.

## Conclusions

Selection against hybrids due to ecological factors, when hybrid phenotypes are ill-adapted to existing ecological niches, has often been considered as being different from selection arising from epistatic (Dobzhansky-Muller) incompatibilities (Thompson et al. 2023). There is a recent push to close this gap between evolutionary ecology and speciation genetics, and to focus more on the role of the environment and ecological adaptation in driving what we traditionally think of as ‘intrinsic’ genetic isolation barriers (Anderson et al. 2023; Thompson et al. 2023). Our study contributes to bridging this gap by synthesizing phenotypic and whole genome data, and demonstrating that divergent ecological adaptation can indeed promote intrinsic reproductive isolation, consistent with predictions of ecological speciation theory. Our population genomics results show that divergent ecological selection, even for relatively few mitotic generations, was accompanied by vast genomic instability in F2 hybrids, including deletions, inversions, and interchromosomal translocations. Our multifaceted approach of combining experimental evolution with sexual microbes, hybridization at increasing time points of divergence, high-throughput phenotyping, and comparative genomics, confirms the ability of ecology to drive reproductive isolation and speciation. Our approach can be further developed to provide a comprehensive empirical framework to test different concepts in speciation research. This includes, for instance, i) testing if the speciation continuum is a ‘continuum of reproductive isolation’ (Stankowski and Ravinet 2021) and whether reproductive isolation barriers accumulate linearly or, as predicted by theory, exponentially (Orr 1995), the ‘missing snowball’ controversy); ii) testing if the ‘speciation hypercube’ captures the complexity of species divergence (Dieckmann et al. 2004; Bolnick et al. 2023), by including more phenotypic and genetic variables, and iii) to better understand the evolution of hybrid fitness during speciation by including additional data sets (e.g. gene interaction networks (Dagilis et al. 2019), which could reconcile the contrasting outcomes of epistatic genetic interactions leading to hybrids with both increased (heterosis) and decreased fitness (hybrid inviability) during different stages of speciation.

## Acknowledgements

We thank Maria de la Paz Celorio Mancera, Viktoria Köppä, and Zebin Zhang for helping with serial transfers and media preparation, and Lorena Ament-Velásquez for bioinformatics advice. We also thank three anonymous reviewers for their constructive comments on an earlier draft. Computation was performed on resources provided by SNIC through Uppsala Multidisciplinary Center for Advanced Computational Science (UPPMAX) under Projects NAISS2023/23-519 (Small Storage) and NAISS2023/22-62 and NAISS2023/22-924 (Small Compute). We acknowledge support from Science for Life Laboratory, the National Genomics Infrastructure, NGI, and Uppmax for providing assistance in massive parallel sequencing and computational infrastructure. This work was supported by Swedish Research Council Project Grants (2022-03427 to R.S. and 2022-03024 to J.W.), a Knut and Alice Wallenberg Foundation Grant (2017.0163 to R.S.), and the Wenner-Gren Foundations (UPD2018-0196, UPD2019-0110 to D.B.).

## Data Accessibility and Benefit-Sharing Section

Phenotypic data are available in Table S1. The raw sequences were deposited in the European Nucleotide Archive, parental populations are available at accession number PRJEB46680, while hybrids are available at accession number PRJEB83568. The code used for analyses and plotting are available at https://github.com/devinbendixsen/EvolvingRI. Benefits from this research accrue from the sharing of our data and results on public databases as described above.

## Author Contributions

Conceived research: Rike Stelkens; Designed research: Rike Stelkens, Ciaran Gilchrist; Performed research: Devin P. Bendixsen, Ciaran Gilchrist, Chloé Haberkorn, Karl Persson; Contributed new reagents or analytical tools: Karl Persson, Jonas Warringer; Data analysis: Devin P. Bendixsen, Chloé Haberkorn, Karl Persson; Writing: Rike Stelkens, Devin P. Bendixsen, Chloé Haberkorn, Ciaran Gilchrist; Funding acquisition: Rike Stelkens, Devin P. Bendixsen, Cecilia Geijer, Jonas Warringer.

## References

Alalam H, Graf FE, Palm M, Abadikhah M, Zackrisson M, Boström J, Fransson A, Hadjineophytou C, Persson L, Stenberg S, et al. 2020. A High-Throughput Method for Screening for Genes Controlling Bacterial Conjugation of Antibiotic Resistance. mSystems 5:e01226–20.

Ament-Velásquez SL, Gilchrist C, Rêgo A, Bendixsen DP, Brice C, Grosse-Sommer JM, Rafati N, Stelkens R. 2022. The Dynamics of Adaptation to Stress from Standing Genetic Variation and de novo Mutations. Molecular Biology and Evolution 39:msac242.

Anderson SAS, López-Fernández H, Weir JT. 2023. Ecology and the Origin of Nonephemeral Species. The American Naturalist 201:619–638.

Anderson SAS, Weir JT. 2022. The role of divergent ecological adaptation during allopatric speciation in vertebrates. Science 378:1214–1218.

Andrews S. 2010. FastQC: A Quality Control tool for High Throughput Sequence Data. Available from: https://www.bioinformatics.babraham.ac.uk/projects/fastqc/

Bautista C, Gagnon-Arsenault I, Utrobina M, Fijarczyk A, Bendixsen DP, Stelkens R, Landry C. 2023. Hybrid adaptation is hampered by Haldane’s sieve. bioRxiv:2023–12.

Bautista C, Marsit S, Landry CR. 2021. Interspecific hybrids show a reduced adaptive potential under DNA damaging conditions. Evolutionary Applications 14:758–769.

Bell G, Gonzalez A. 2011. Adaptation and Evolutionary Rescue in Metapopulations Experiencing Environmental Deterioration. Science 332:1327–1330.

Belleau P, Deschênes A, Beyaz S, Tuveson DA, Krasnitz A. 2021. CNVMetrics package: Quantifying similarity between copy number profiles. F1000Research [Internet] 10. Available from: https://f1000research.com/slides/10-737

Bendixsen DP, Gettle N, Gilchrist C, Zhang Z, Stelkens R. 2021. Genomic Evidence of an Ancient East Asian Divergence Event in Wild Saccharomyces cerevisiae. Genome Biology and Evolution 13:evab001.

Bendixsen DP, Peris D, Stelkens R. 2021. Patterns of Genomic Instability in Interspecific Yeast Hybrids With Diverse Ancestries. Frontiers in Fungal Biology [Internet] 2. Available from: https://www.frontiersin.org/article/10.3389/ffunb.2021.742894

Bernardes JP, Stelkens RB, Greig D. 2017. Heterosis in hybrids within and between yeast species. J Evol Biol 30:538–548.

Boeva V, Popova T, Bleakley K, Chiche P, Cappo J, Schleiermacher G, Janoueix-Lerosey I, Delattre O, Barillot E. 2012. Control-FREEC: a tool for assessing copy number and allelic content using next-generation sequencing data. Bioinformatics 28:423–425.

Boeva V, Zinovyev A, Bleakley K, Vert J-P, Janoueix-Lerosey I, Delattre O, Barillot E. 2011. Control-free calling of copy number alterations in deep-sequencing data using GC-content normalization. Bioinformatics 27:268–269.

Bolger AM, Lohse M, Usadel B. 2014. Trimmomatic: a flexible trimmer for Illumina sequence data. Bioinformatics 30:2114–2120.

Bolnick DI, Hund AK, Nosil P, Peng F, Ravinet M, Stankowski S, Subramanian S, Wolf JBW, Yukilevich R. 2023. A multivariate view of the speciation continuum. Evolution 77:318–328.

Bozdag GO, Ono J, Denton JA, Karakoc E, Hunter N, Leu J-Y, Greig D. 2021. Breaking a species barrier by enabling hybrid recombination. Current Biology 31:R180–R181.

Brice C, Zhang Z, Bendixsen D, Stelkens R. 2021. Hybridization Outcomes Have Strong Genomic and Environmental Contingencies. The American Naturalist 198:E53–E67.

Cervantes S, Kesälahti R, Kumpula TA, Mattila TM, Helanterä H, Pyhäjärvi T. 2023. Strong Purifying Selection in Haploid Tissue–Specific Genes of Scots Pine Supports the Masking Theory. Molecular Biology and Evolution 40:msad183.

Chang S-L, Lai H-Y, Tung S-Y, Leu J-Y. 2013. Dynamic Large-Scale Chromosomal Rearrangements Fuel Rapid Adaptation in Yeast Populations. PLOS Genetics 9:e1003232.

Chen X, Schulz-Trieglaff O, Shaw R, Barnes B, Schlesinger F, Källberg M, Cox AJ, Kruglyak S, Saunders CT. 2016. Manta: rapid detection of structural variants and indels for germline and cancer sequencing applications. Bioinformatics 32:1220–1222.

Cingolani P, Platts A, Wang LL, Coon M, Nguyen T, Wang L, Land SJ, Lu X, Ruden DM. 2012. A program for annotating and predicting the effects of single nucleotide polymorphisms, SnpEff: SNPs in the genome of Drosophila melanogaster strain w1118; iso-2; iso-3. Fly (Austin) 6:80–92.

Dagilis AJ, Kirkpatrick M, Bolnick DI. 2019. The evolution of hybrid fitness during speciation. PLOS Genetics 15:e1008125.

Danecek P, Auton A, Abecasis G, Albers CA, Banks E, DePristo MA, Handsaker RE, Lunter G, Marth GT, Sherry ST, et al. 2011. The variant call format and VCFtools. Bioinformatics 27:2156–2158.

Danecek P, Bonfield JK, Liddle J, Marshall J, Ohan V, Pollard MO, Whitwham A, Keane T, McCarthy SA, Davies RM, et al. 2021. Twelve years of SAMtools and BCFtools. GigaScience 10:giab008.

Deschênes A, Belleau P, Tuveson DA, Krasnitz A. 2022. Quantifying similarity between copy number profiles with CNVMetrics package. F1000Research [Internet] 11. Available from: https://f1000research.com/posters/11-816

Dettman JR, Anderson JB, Kohn LM. 2008. Divergent adaptation promotes reproductive isolation among experimental populations of the filamentous fungus Neurospora. BMC Evol Biol 8:35.

Dettman JR, Sirjusingh C, Kohn LM, Anderson JB. 2007. Incipient speciation by divergent adaptation and antagonistic epistasis in yeast. Nature 447:585–588.

deVicente MC Tanksley SD. 1993. Qtl Analysis of Transgressive Segregation in an Interspecific Tomato Cross. Genetics 134:585–596.

Dieckmann U, Doebeli M, Metz JAJ, Tautz D eds. 2004. Adaptive Speciation. Cambridge: Cambridge University Press Available from: https://www.cambridge.org/core/books/adaptive-speciation/A2413E0B1E5DDAB5C5848F0666468A35

Dodd DMB. 1989. Reproductive Isolation as a Consequence of Adaptive Divergence in Drosophila pseudoobscura. Evolution 43:1308–1311.

Eberlein C, Hénault M, Fijarczyk A, Charron G, Bouvier M, Kohn LM, Anderson JB, Landry CR. 2019. Hybridization is a recurrent evolutionary stimulus in wild yeast speciation. Nat Commun 10:923.

Filchak KE, Roethele JB, Feder JL. 2000. Natural selection and sympatric divergence in the apple maggot Rhagoletis pomonella. Nature 407:739–742.

Forejt J, Jansa P. 2023. Meiotic Recognition of Evolutionarily Diverged Homologs: Chromosomal Hybrid Sterility Revisited. Molecular Biology and Evolution 40:msad083.

Funk DJ, Nosil P, Etges WJ. 2006. Ecological divergence exhibits consistently positive associations with reproductive isolation across disparate taxa. Proceedings of the National Academy of Sciences 103:3209–3213.

Gallone B, Steensels J, Mertens S, Dzialo MC, Gordon JL, Wauters R, Theßeling FA, Bellinazzo F, Saels V, Herrera-Malaver B, et al. 2019. Interspecific hybridization facilitates niche adaptation in beer yeast. Nat Ecol Evol 3:1562–1575.

Garrison E, Marth G. 2012. Haplotype-based variant detection from short-read sequencing. *arXiv* 1207.

Gautier M, Vitalis R, Flori L, Estoup A. 2022. f-Statistics estimation and admixture graph construction with Pool-Seq or allele count data using the R package poolfstat. Mol Ecol Resour 22:1394–1416.

Gerstein AC, Cleathero LA, Mandegar MA, Otto SP. 2011. Haploids adapt faster than diploids across a range of environments. Journal of Evolutionary Biology 24:531–540.

Gilchrist C, Stelkens R. 2019. Aneuploidy in yeast: Segregation error or adaptation mechanism? Yeast 36:525–539.

Good BH, McDonald MJ, Barrick JE, Lenski RE, Desai MM. 2017. The dynamics of molecular evolution over 60,000 generations. Nature 551:45–50.

Gorter FA, Derks MF, van den Heuvel J, Aarts MG, Zwaan BJ, de Ridder D, de Visser JAG. 2017. Genomics of adaptation depends on the rate of environmental change in experimental yeast populations. Molecular Biology and Evolution 34:2613–2626.

Gorter FA, Scanlan PD, Buckling A. 2016. Adaptation to abiotic conditions drives local adaptation in bacteria and viruses coevolving in heterogeneous environments. Biol. Lett. 12:20150879.

Grant RB, Grant PR. 2003. What Darwin’s Finches Can Teach Us about the Evolutionary Origin and Regulation of Biodiversity. BioScience 53:965–975.

Greig D, Louis EJ, Borts RH, Travisano M. 2002. Hybrid Speciation in Experimental Populations of Yeast. Science 298:1773–1775.

Härer A, Bolnick DI, Rennison DJ. 2021. The genomic signature of ecological divergence along the benthic-limnetic axis in allopatric and sympatric threespine stickleback. Molecular Ecology 30:451–463.

Hatfield T, Schluter D. 1999. ECOLOGICAL SPECIATION IN STICKLEBACKS: ENVIRONMENT-DEPENDENT HYBRID FITNESS. Evolution 53:866–873.

Hendry AP, Nosil P, Rieseberg LH. 2007. The speed of ecological speciation. Functional Ecology 21:455– 464.

Hivert V, Leblois R, Petit EJ, Gautier M, Vitalis R. 2018. Measuring Genetic Differentiation from Pool-seq Data. Genetics 210:315–330.

Hunter N, Chambers SR, Louis EJ, Borts RH. 1996. The mismatch repair system contributes to meiotic sterility in an interspecific yeast hybrid. EMBO J 15:1726–1733.

Irestedt M, Müller I, Thörn F, Joseph L, Nylander J, Guinet B, Valk TV der, Jønsson K. Reticulate and hybrid speciation is promoted by environmental instability in an Indo-Pacific species complex of whistlers (Aves: Pachycephala). Available from: https://www.authorea.com/doi/full/10.22541/au.173053090.08198521?commit=f442f52dffaefd5b78602e94558c1591a8dff4e5

Jarrett BJM, Downing PA, Svensson EI. 2025. Meta-analysis reveals that phenotypic plasticity and divergent selection promote reproductive isolation during incipient speciation. Nat Ecol Evol 9:833–844.

Johnson MS, Desai MM. 2022. Mutational robustness changes during long-term adaptation in laboratory budding yeast populations. Cooper VS, Walczak AM, Miller C, editors. eLife 11:e76491.

Kirkpatrick M, Ravigné V. 2002. Speciation by natural and sexual selection: models and experiments. Am Nat 159 Suppl 3:S22–35.

Kosheleva K, Desai MM. 2018. Recombination Alters the Dynamics of Adaptation on Standing Variation in Laboratory Yeast Populations. Mol Biol Evol 35:180–201.

Lenski RE, Rose MR, Simpson SC, Tadler SC. 1991. Long-Term Experimental Evolution in Escherichia coli. I. Adaptation and Divergence During 2,000 Generations. The American Naturalist 138:1315–1341.

Leu J-Y, Chang S-L, Chao J-C, Woods LC, McDonald MJ. 2020. Sex alters molecular evolution in diploid experimental populations of S. cerevisiae. Nat Ecol Evol 4:453–460.

Li H, Durbin R. 2009. Fast and accurate short read alignment with Burrows–Wheeler transform. Bioinformatics 25:1754–1760.

Li J, Vázquez-García I, Persson K, González A, Yue J-X, Barré B, Hall MN, Long A, Warringer J, Mustonen V, et al. 2019. Shared Molecular Targets Confer Resistance over Short and Long Evolutionary Timescales. Molecular Biology and Evolution 36:691–708.

Liu Y, Ma G, Gao Z, Li J, Wang J, Zhu X, Ma R, Yang J, Zhou Y, Hu K, et al. 2022. Global chromosome rearrangement induced by CRISPR-Cas9 reshapes the genome and transcriptome of human cells. Nucleic Acids Research 50:3456–3474.

Louder MIM, Justen H, Kimmitt AA, Lawley KS, Turner LM, Dickman JD, Delmore KE. 2024. Gene regulation and speciation in a migratory divide between songbirds. Nat Commun 15:98.

Mayr E. 1947. Ecological Factors in Speciation. Evolution 1:263–288.

McDonald MJ, Rice DP, Desai MM. 2016. Sex speeds adaptation by altering the dynamics of molecular evolution. Nature 531:233–236.

McKinnon JS, Mori S, Blackman BK, David L, Kingsley DM, Jamieson L, Chou J, Schluter D. 2004. Evidence for ecology’s role in speciation. Nature 429:294–298.

Meyer JR, Dobias DT, Medina SJ, Servilio L, Gupta A, Lenski RE. 2016. Ecological speciation of bacteriophage lambda in allopatry and sympatry. Science 354:1301–1304.

Michel AP, Sim S, Powell THQ, Taylor MS, Nosil P, Feder JL. 2010. Widespread genomic divergence during sympatric speciation. Proceedings of the National Academy of Sciences 107:9724–9729.

Muenzner J, Trébulle P, Agostini F, Zauber H, Messner CB, Steger M, Kilian C, Lau K, Barthel N, Lehmann A, et al. 2024. Natural proteome diversity links aneuploidy tolerance to protein turnover. Nature 630:149–157.

Mukherjee V, Lind U, St. Onge RP, Blomberg A, Nygård Y. 2021. A CRISPR Interference Screen of Essential Genes Reveals that Proteasome Regulation Dictates Acetic Acid Tolerance in Saccharomyces cerevisiae. mSystems 6:10.1128/msystems.00418-21.

Müller I, Thörn F, Rajan S, Olsen R-A, Ericson P, Peona V, Smith B, Maiah G, Koane B, Iova B, et al. Ephemeral speciation in a New Guinean honeyeater complex (Aves: Melidectes). Available from: https://www.authorea.com/doi/full/10.22541/au.173639442.28305532?commit=c6fffa9cc0ca71d3e44f2de72f5001a07ea2ab20

Munar-Delgado G, Pulido F, Edelaar P. 2024. Performance-based habitat choice can drive rapid adaptive divergence and reproductive isolation. Current Biology 34:5564–5569.e4.

Nosil P. 2012. Ecological Speciation. Oxford University Press Available from: https://academic.oup.com/book/5499

Nosil P, Crespi BJ. 2006. Experimental evidence that predation promotes divergence in adaptive radiation. Proceedings of the National Academy of Sciences 103:9090–9095.

Nosil P, Feder JL. 2012. Genomic divergence during speciation: causes and consequences. Philosophical Transactions of the Royal Society B: Biological Sciences 367:332–342.

Nosil P, Schluter D. 2011. The genes underlying the process of speciation. Trends Ecol Evol 26:160–167.

Okonechnikov K, Conesa A, García-Alcalde F. 2016. Qualimap 2: advanced multi-sample quality control for high-throughput sequencing data. Bioinformatics 32:292–294.

Ono J, Gerstein AC, Otto SP. 2017. Widespread Genetic Incompatibilities between First-Step Mutations during Parallel Adaptation of Saccharomyces cerevisiae to a Common Environment. PLOS Biology 15:e1002591.

Orr HA. 1995. The population genetics of speciation: the evolution of hybrid incompatibilities. Genetics 139:1805–1813.

Persson K, Onyema VO, Nwafor IP, Peri KVR, Otti C, Nnaemeka P, Onyishi C, Okoye S, Moneke A, Amadi O, et al. 2024. Lactose-assimilating yeasts with high fatty acid accumulation uncovered by untargeted bioprospecting. Applied and Environmental Microbiology 0:e01615–24.

Persson K, Stenberg S, Tamás MJ, Warringer J. 2022. Adaptation of the yeast gene knockout collection is near-perfectly predicted by fitness and diminishing return epistasis. G3 Genes|Genomes|Genetics 12:jkac240.

Peter J, De Chiara M, Friedrich A, Yue J-X, Pflieger D, Bergström A, Sigwalt A, Barre B, Freel K, Llored A, et al. 2018. Genome evolution across 1,011 Saccharomyces cerevisiae isolates. Nature 556:339– 344.

R Core Team. 2023. R: A Language and Environment for Statistical Computing. Available from: https://www.R-project.org/

Rainey PB, Travisano M. 1998. Adaptive radiation in a heterogeneous environment. Nature 394:69–72.

Rice WR, Hostert EE. 1993. Laboratory Experiments on Speciation: What Have We Learned in 40 Years? Evolution 47:1637–1653.

Rogers DW, McConnell E, Ono J, Greig D. 2018. Spore-autonomous fluorescent protein expression identifies meiotic chromosome mis-segregation as the principal cause of hybrid sterility in yeast. PLOS Biology 16:e2005066.

Rundle HD, Nagel L, Wenrick Boughman J, Schluter D. 2000. Natural selection and parallel speciation in sympatric sticklebacks. Science 287:306–308.

Sawada A, Iwasaki T, Akatani K, Takagi M. 2022. Mate choice for body size leads to size assortative mating in the Ryukyu Scops Owl Otus elegans. Ecology and Evolution 12:e9578.

Schluter D. 2001. Ecology and the origin of species. Trends Ecol Evol 16:372–380.

Schluter D. 2009. Evidence for Ecological Speciation and Its Alternative. Science 323:737–741.

Scopel EFC, Hose J, Bensasson D, Gasch AP. 2021. Genetic variation in aneuploidy prevalence and tolerance across Saccharomyces cerevisiae lineages. Genetics 217:iyab015.

Shapira R, Levy T, Shaked S, Fridman E, David L. 2014. Extensive heterosis in growth of yeast hybrids is explained by a combination of genetic models. Heredity (Edinb) 113:316–326.

Sparks MM, Kraft JC, Blackstone KMS, McNickle GG, Christie MR. 2022. Large genetic divergence underpins cryptic local adaptation across ecological and evolutionary gradients. Proceedings of the Royal Society B: Biological Sciences 289:20221472.

Stankowski S, Ravinet M. 2021. Defining the speciation continuum. Evolution 75:1256–1273.

Stelkens R. 2024. A microbial perspective on speciation. Evolutionary Journal of the Linnean Society 3:kzae023.

Stelkens RB, Brockhurst MA, Hurst GDD, Miller EL, Greig D. 2014. The effect of hybrid transgression on environmental tolerance in experimental yeast crosses. J. Evol. Biol. 27:2507–2519.

Stenberg S, Li J, Gjuvsland AB, Persson K, Demitz-Helin E, González Peña C, Yue J-X, Gilchrist C, Ärengård T, Ghiaci P, et al. 2022. Genetically controlled mtDNA deletions prevent ROS damage by arresting oxidative phosphorylation. Gruber J, Tyler JK, Toledano MB, editors. eLife 11:e76095.

Struck TH, Feder JL, Bendiksby M, Birkeland S, Cerca J, Gusarov VI, Kistenich S, Larsson K-H, Liow LH, Nowak MD, et al. 2018. Finding Evolutionary Processes Hidden in Cryptic Species. Trends in Ecology & Evolution 33:153–163.

Tange O. 2011. Gnu parallel-the command-line power tool. Usenix Mag 36:42.

Thompson KA, Brandvain Y, Coughlan JM, Delmore KE, Justen H, Linnen CR, Ortiz-Barrientos D, Rushworth CA, Schneemann H, Schumer M, et al. 2023. The Ecology of Hybrid Incompatibilities. Cold Spring Harb Perspect Biol: a041440.

Tusso S, Nieuwenhuis BPS, Weissensteiner B, Immler S, Wolf JBW. 2021. Experimental evolution of adaptive divergence under varying degrees of gene flow. Nat Ecol Evol 5:338–349.

Villa SM, Altuna JC, Ruff JS, Beach AB, Mulvey LI, Poole EJ, Campbell HE, Johnson KP, Shapiro MD, Bush SE, et al. 2019. Rapid experimental evolution of reproductive isolation from a single natural population. Proceedings of the National Academy of Sciences 116:13440–13445.

Westram AM, Stankowski S, Surendranadh P, Barton N. 2022. What is reproductive isolation? Journal of Evolutionary Biology 35:1143–1164.

Zackrisson M, Hallin J, Ottosson L-G, Dahl P, Fernandez-Parada E, Ländström E, Fernandez-Ricaud L, Kaferle P, Skyman A, Stenberg S, et al. 2016. Scan-o-matic: High-Resolution Microbial Phenomics at a Massive Scale. G3 Genes|Genomes|Genetics 6:3003–3014.

Zhang Z, Bendixsen DP, Janzen T, Nolte AW, Greig D, Stelkens R. 2020. Recombining Your Way Out of Trouble: The Genetic Architecture of Hybrid Fitness under Environmental Stress. Molecular Biology and Evolution 37:167–182.

