## Supplementary Figures for "Reproductive isolation due to divergent ecological selection is accompanied by vast genomic instability in experimentally evolved yeast populations"

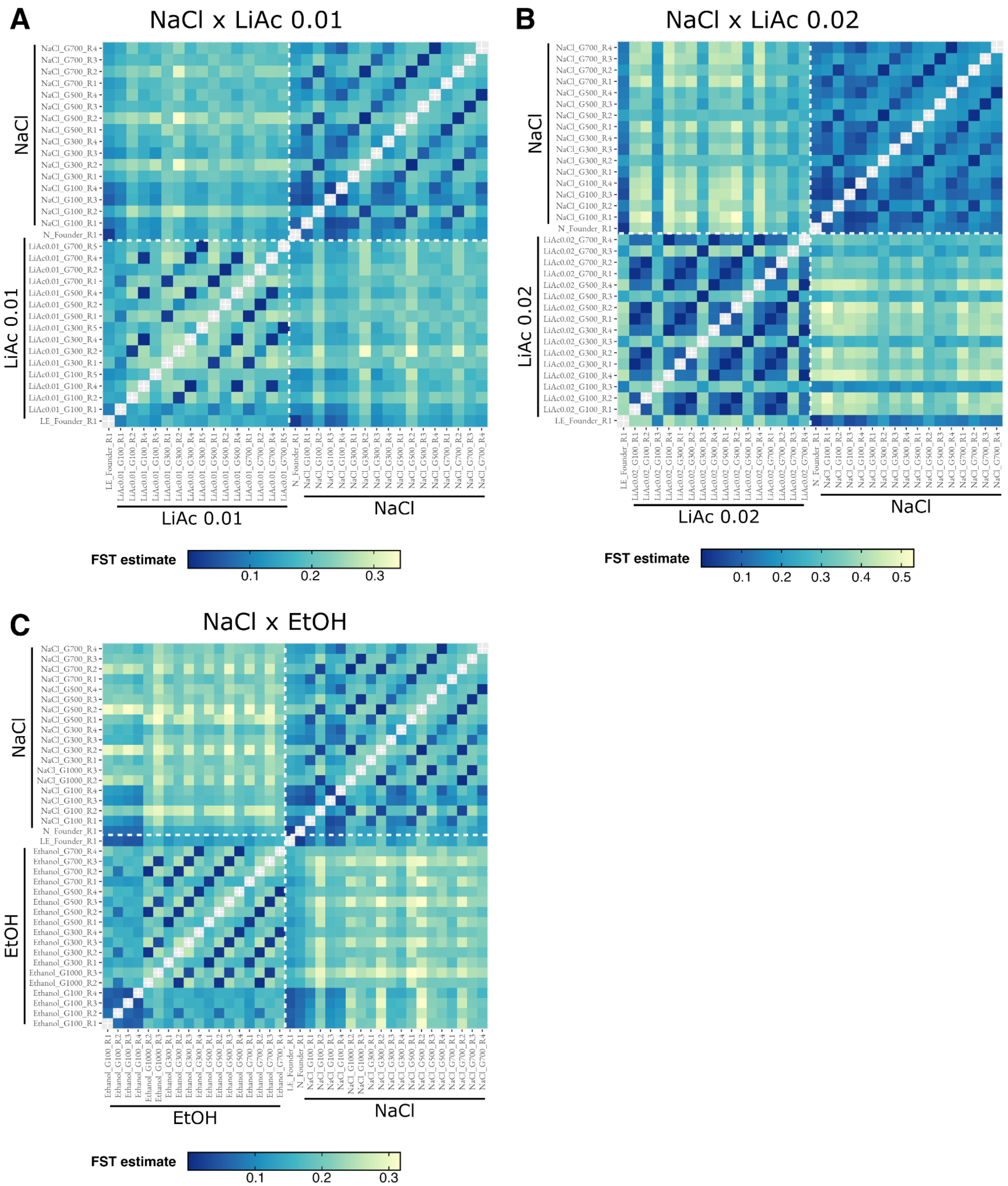


**Figure S1: Pairwise $F_{ST}$ estimates over the course of experimental evolution.** $F_{ST}$ estimates between replicate (divergent- and parallel-selected) parental populations and timepoints for **A)** NaCl x LiAc0.01, **B)** NaCl x LiAc0.02 and **C)** NaCl x Ethanol. The colour scale indicates pairwise $F_{ST}$ estimate with dark blue indicating smaller $F_{ST}$ suggesting less population differentiation, and lighter yellow indicating higher $F_{ST}$ suggesting greater population differentiation.

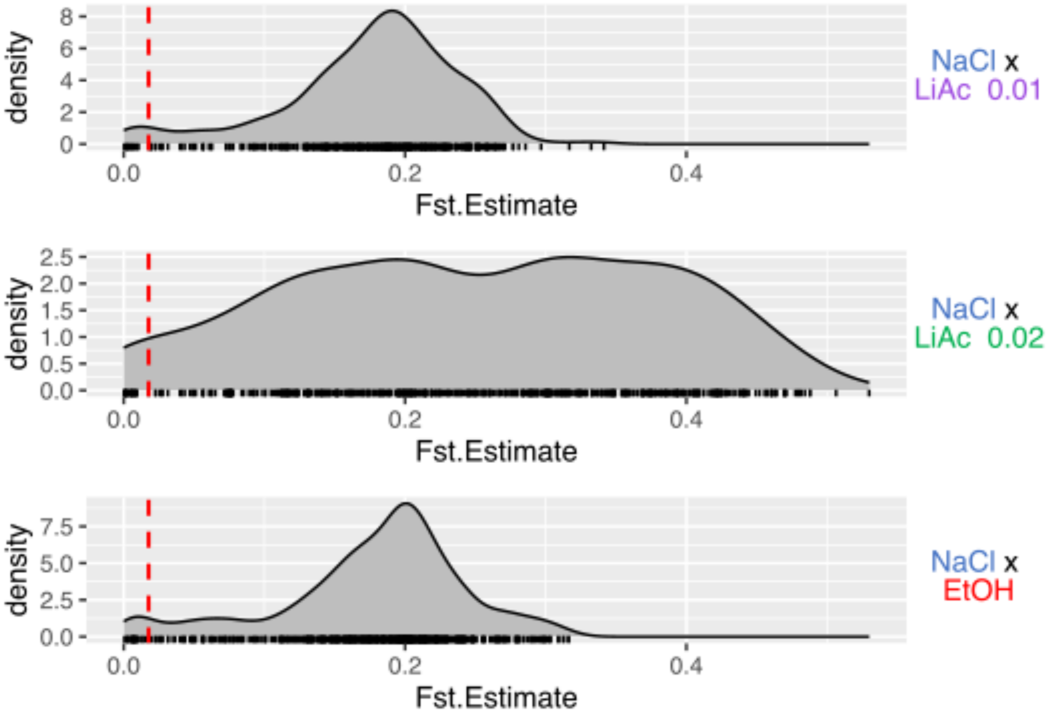


**Figure S2: The distribution of pairwise $F_{ST}$ estimates over the course of experimental evolution.** Each density plot indicates the distribution of pairwise values between replicates (divergent and parallel-selected) parental populations. The dashed red line indicates the estimate between the two founder populations (N Founder and LE Founder).

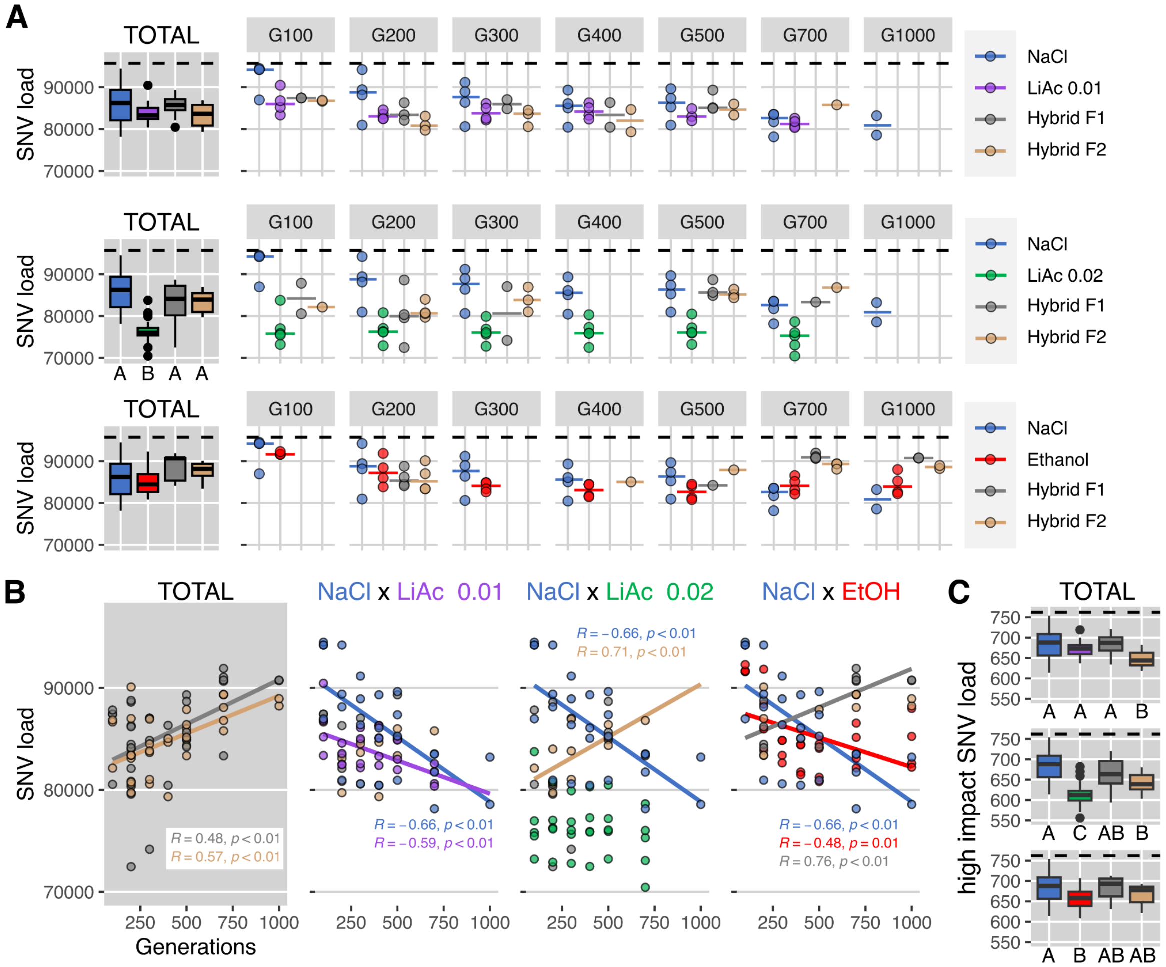


**Figure S3: The single nucleotide variant mutational landscape of hybridization** at seven time points of experimental evolution. **A)** The SNV mutational load consists of the number of mutational variants and indels (insertions and deletions) within a population. The median SNV mutational load of populations is indicated by the colored solid line. Gray plots show mutational load averaged across all sampled timepoints (TOTAL) for the three types of hybrid crosses. Other plots show data from crosses made at 7 timepoints of increasing parental divergence (number of generations). The mean mutational load of the founder population is indicated by the dashed black line. Boxplots indicate the first and third quartiles with whiskers extending to the furthest point not exceeding 1.5 x the interquartile range and outliers beyond this are shown as dots. The solid line indicates the median value. Letters indicate statistical significance using Kruskal-Wallis and pairwise Wilcoxon post hoc tests with Bonferroni corrections. **B)** Linear regression (Pearson R) of SNV mutational load at seven time points (n of generations) of experimental evolution. Only significant linear relationships are shown (p ≤ 0.05). The gray plot includes F1 and F2 hybrids across all crosses. **C)** SNV mutational load after filtering for high impact mutations on gene function using SnpEff v5.2 (Cingolani et al. 2012). The mutational load is averaged across all timepoints (TOTAL) for each hybrid cross. The mean mutational load of the founder population is indicated by the dashed black line. Different letters (A, B and C) indicate statistically significant differences using Kruskal-Wallis and pairwise Wilcoxon post hoc tests with Bonferroni corrections.

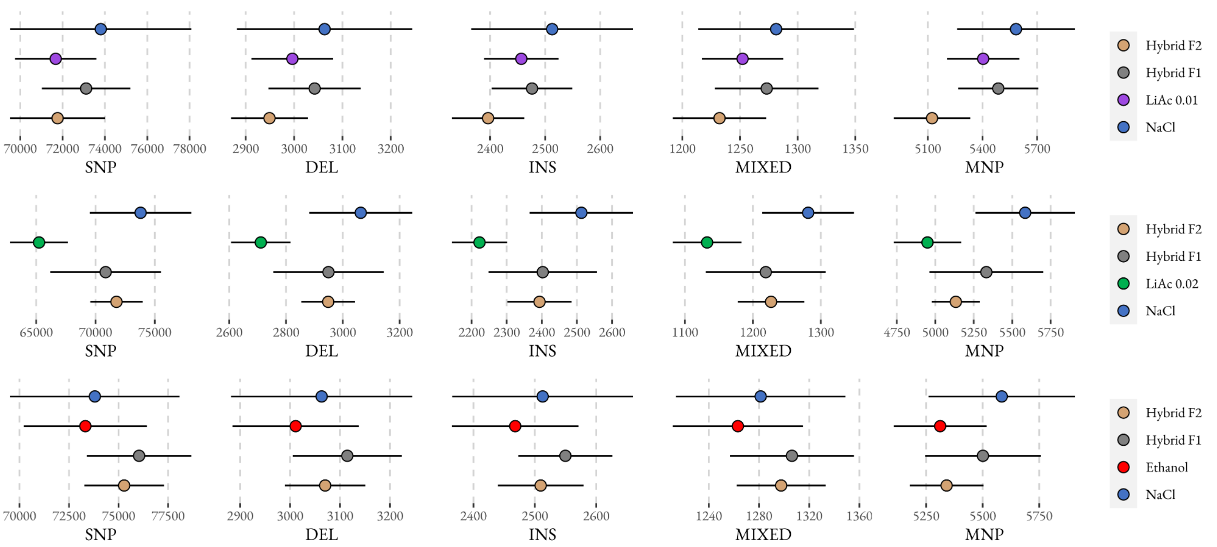


**Figure S4 The SNV mutational landscape sorted by mutational type.** The overall mean and standard deviation of each mutation type (SNP = single nucleotide polymorphism, DEL = deletion, INS = insertion, MNP = multiple nucleotide polymorphism, MIXED = multiple nucleotide polymorphism + indels) in parental, F1 and F2 hybrid populations across all sampled timepoints.

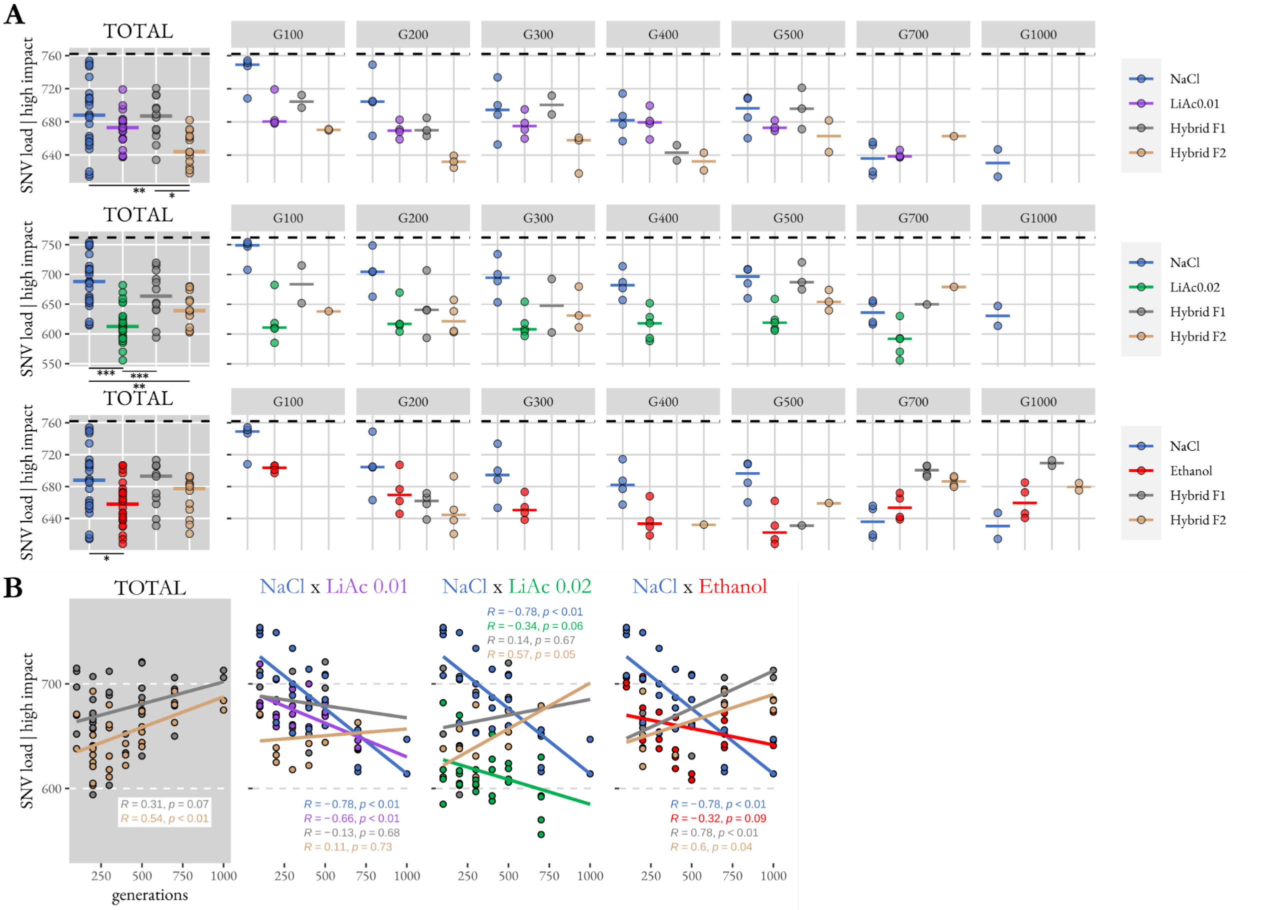


**Figure S5 The high impact SNV mutational landscape of hybridization. A)** The SNV mutational load consists of mutational variants and indels (insertions and deletions) that have been estimated to have a high impact on gene function by SnpEff v5.2 (Cingolani et al. 2012). The mutational load is shown encompassing all timepoints (TOTAL) for a given hybrid cross, as well as separately according to the generational divergence of each cross from the initial founder population. The mean mutational load of the two founding populations is indicated by the dashed black line. **B)** Linear regression (Pearson R) between SNV mutational load and parental divergence in terms of generational time. Only significant linear relationships are shown ($p \leq 0.05$).

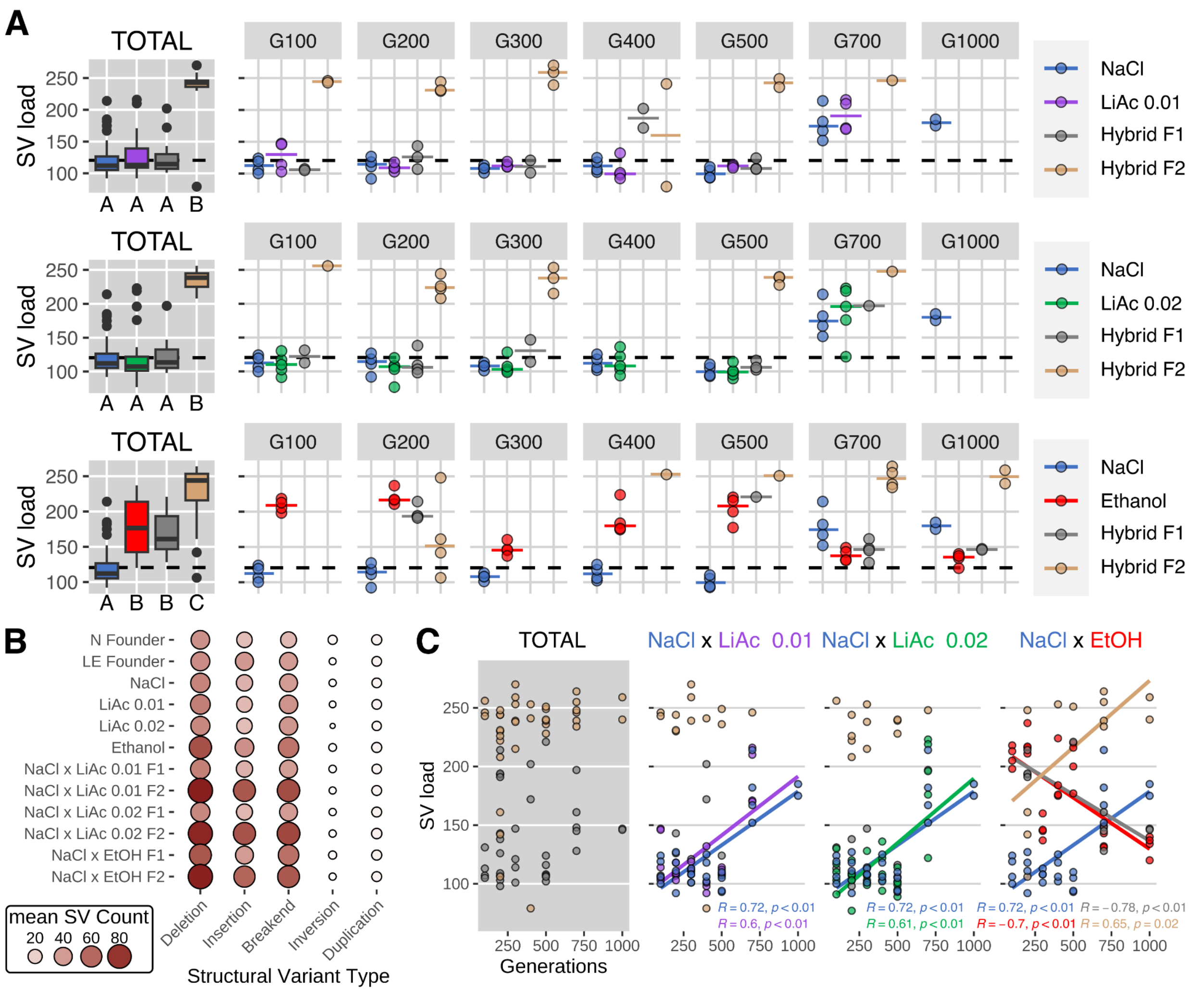


**Figure S6: The structural variant mutational landscape of hybridization** at seven time points of experimental evolution. **A)** Structural variant (SV) mutational load consists of the number of deletions, insertions, inversions, tandem duplications and interchromosomal translocations (> 8bp) detected in the population. Gray plots show SV mutational load averaged across all sampled timepoints (TOTAL) for the three types of hybrid crosses. Other plots show data from crosses made at 7 timepoints of increasing parental divergence (number of generations). The mean mutational load of the founder population is indicated by the dashed black line. Boxplots indicate the first and third quartiles with whiskers extending to the furthest point not exceeding 1.5 x the interquartile range and outliers beyond this are shown as dots. The solid line indicates the median value. Different letters (A, B and C) indicate statistically significant differences using Kruskal-Wallis and pairwise Wilcoxon post hoc tests with Bonferroni corrections. **B)** Mean structural variant count of each type of structural variant (deletion, insertion, breakend, inversion and duplication) for each environment and cross. Node color and size indicates the mean count of each variant type across all timepoints. **C)** Linear regression (Pearson R) between SV mutational load and increasing timepoints of parental divergence (number of generations). Only significant linear relationships are shown ($p \leq 0.05$). The gray plot includes F1 and F2 hybrids from all crosses.

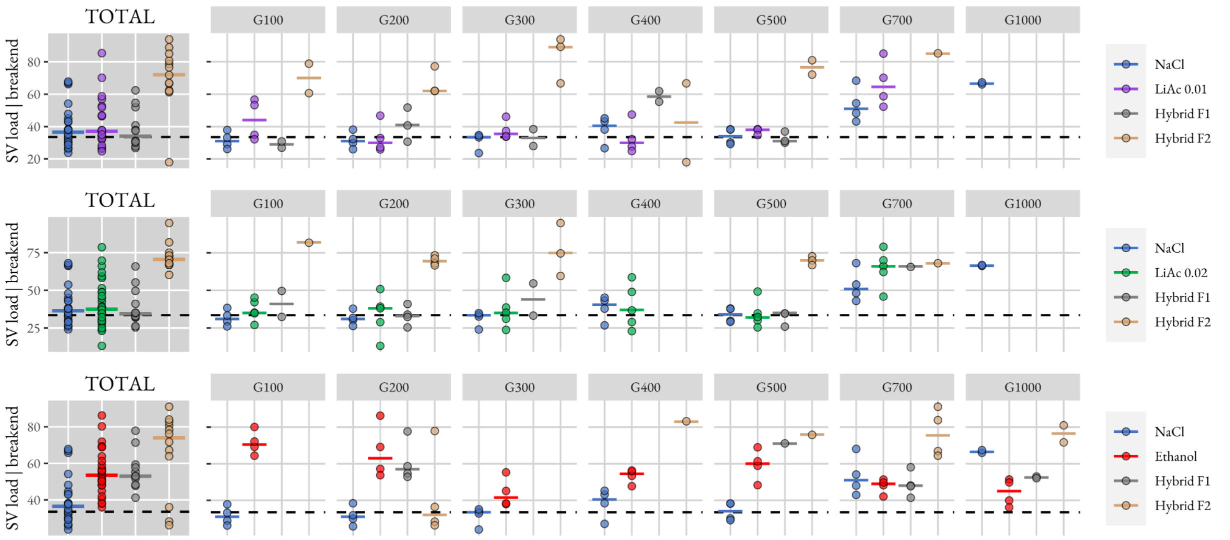


**Figure S7: The distribution of breakend structural variants.** Gray plots show SV breakend load averaged across all sampled timepoints (TOTAL) for the three types of hybrid crosses. Other plots show data from crosses made at 7 timepoints of increasing parental divergence (number of generations). The mean mutational load of the founder population is indicated by the dashed black line.

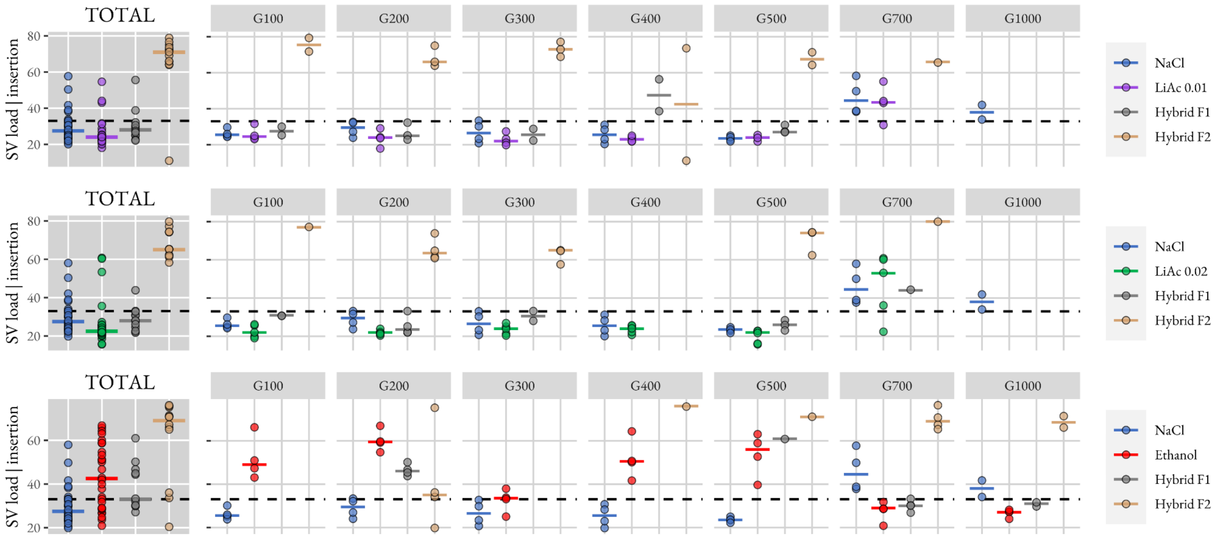


**Figure S8: The distribution of insertion structural variants.** Gray plots show SV insertion load averaged across all sampled timepoints (TOTAL) for the three types of hybrid crosses. Other plots show data from crosses made at 7 timepoints of increasing parental divergence (number of generations). The mean mutational load of the founder population is indicated by the dashed black line.

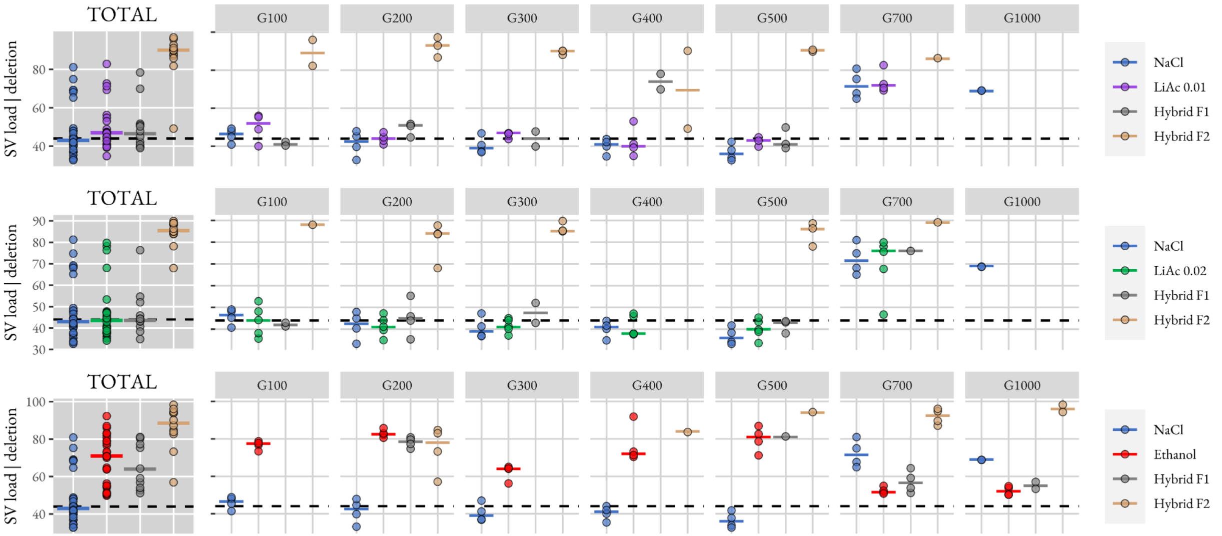


**Figure S9: The distribution of deletion structural variants.** Gray plots show SV deletion load averaged across all sampled timepoints (TOTAL) for the three types of hybrid crosses. Other plots show data from crosses made at 7 timepoints of increasing parental divergence (number of generations). The mean mutational load of the founder population is indicated by the dashed black line.

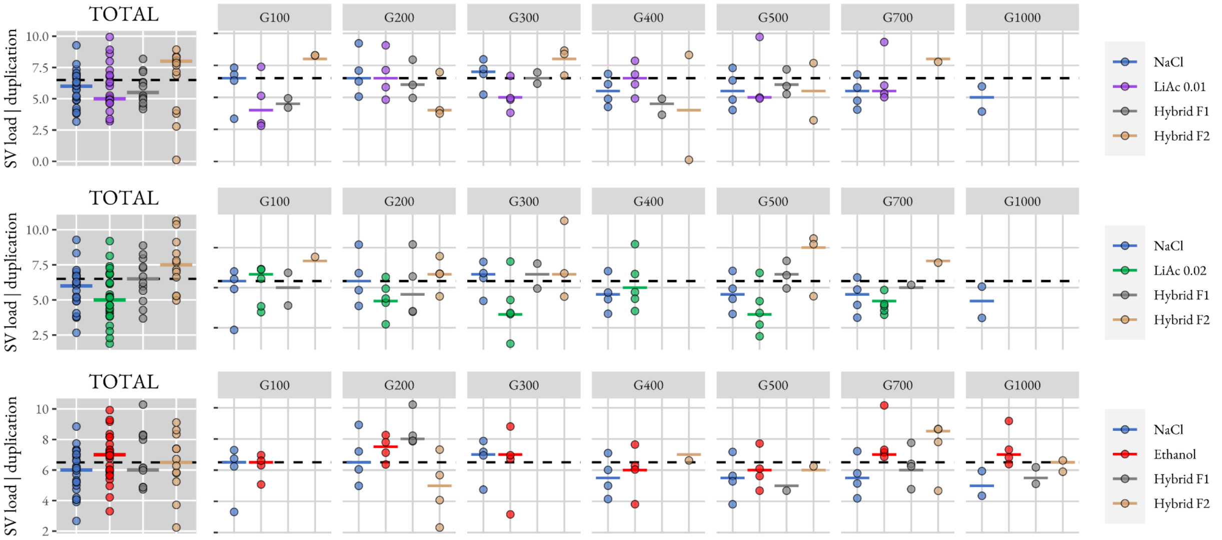


**Figure S10: The distribution of duplication structural variants.** Gray plots show SV duplication load averaged across all sampled timepoints (TOTAL) for the three types of hybrid crosses. Other plots show data from crosses made at 7 timepoints of increasing parental divergence (number of generations). The mean mutational load of the founder population is indicated by the dashed black line.

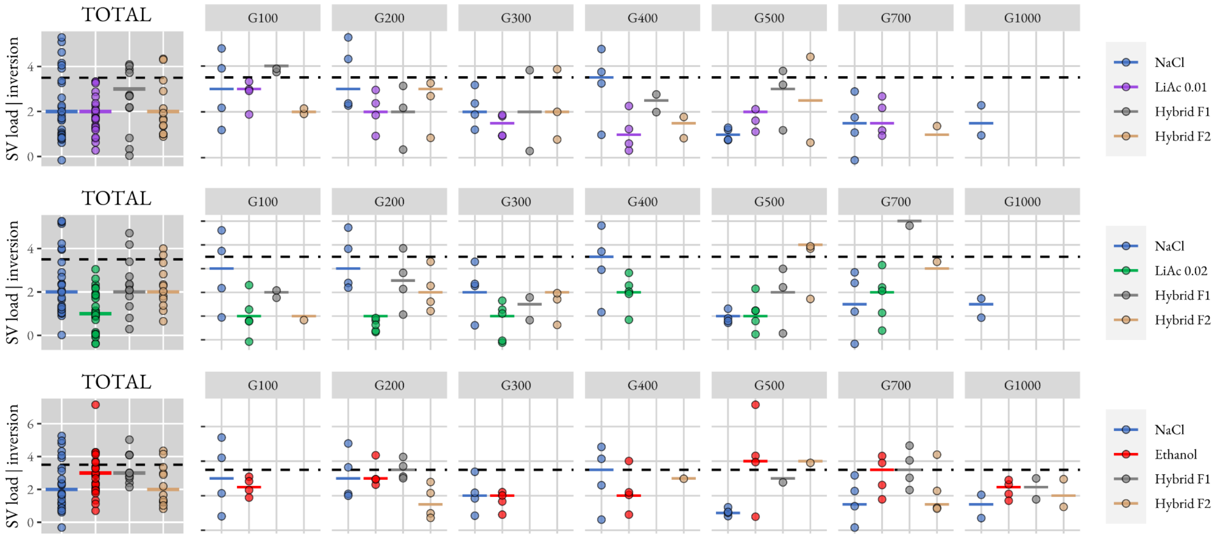


**Figure S11: The distribution of inversion structural variants.** Gray plots show SV inversion load averaged across all sampled timepoints (TOTAL) for the three types of hybrid crosses. Other plots show data from crosses made at 7 timepoints of increasing parental divergence (number of generations). The mean mutational load of the founder population is indicated by the dashed black line.

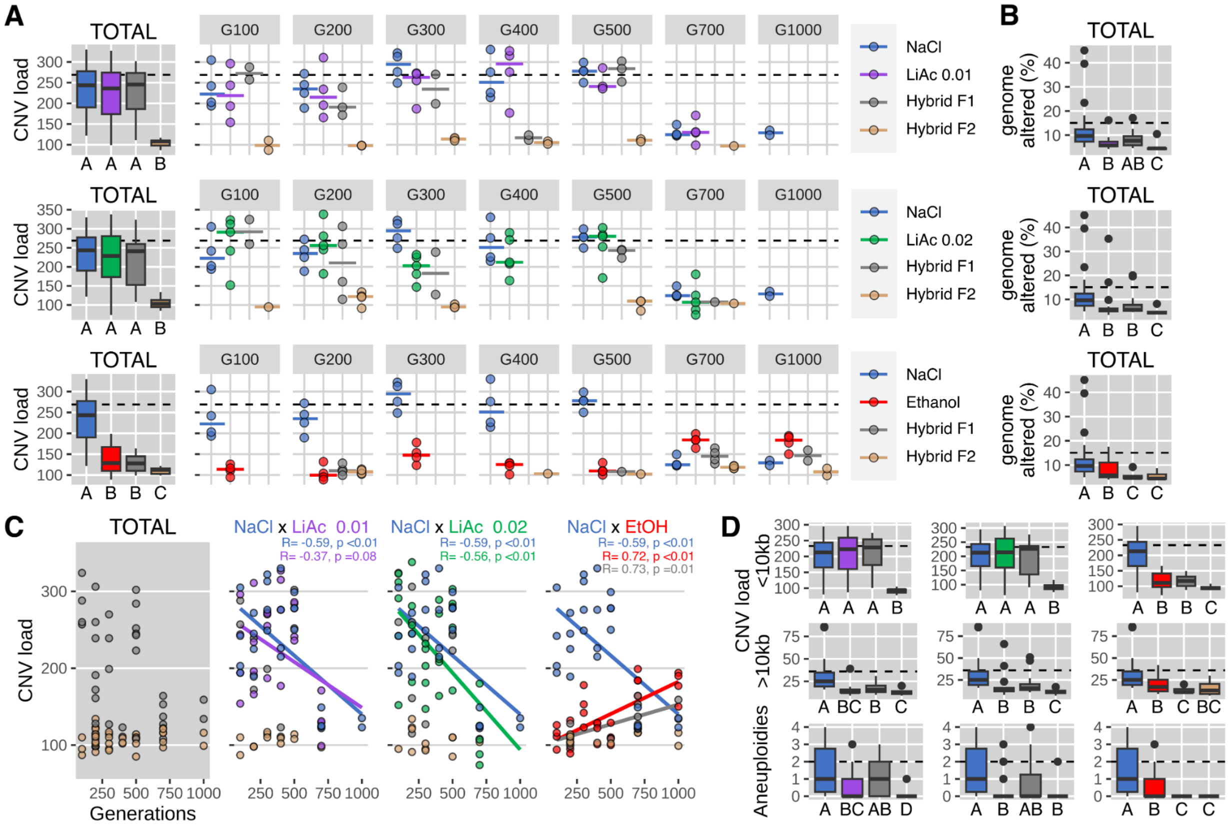


**Figure S12: The copy number variant mutational landscape of hybridization** at seven time points of experimental evolution. **A)** Copy number variant (CNV) mutational load consisting of amplifications, deletions and aneuploidies. Gray plots show CNV mutational load averaged across all sampled timepoints (TOTAL) for the three types of hybrid crosses. Other plots show data from crosses made at 7 timepoints of increasing parental divergence (number of generations). The mean mutational load of the founder population is indicated by the dashed black line. Boxplots indicate the first and third quartiles with whiskers extending to the furthest point not exceeding 1.5 x the interquartile range and outliers beyond this are shown as dots. The solid line indicates the median value. Different letters (A, B and C) indicate statistically significant differences using Kruskal-Wallis and pairwise Wilcoxon post hoc tests with Bonferroni corrections. **B)** The fraction of the genome altered (%) across all sampled timepoints (TOTAL) for the three types of hybrid crosses. Colors, boxplots and statistical significance are consistent with panel A. **C)** Linear regression (Pearson R) between CNV mutational load and increasing timepoints of parental divergence (number of generations). Only significant linear relationships are shown ($p \leq 0.05$). The gray plot includes F1 and F2 hybrids from all crosses. **D)** The distribution of large (>10kb) CNVs, small (< 10kb) CNVs and chromosome aneuploidy (> 60% amplified or deleted) across all sampled timepoints for the three types of hybrid crosses. Colors, boxplots, and statistical significance are consistent with panel A.

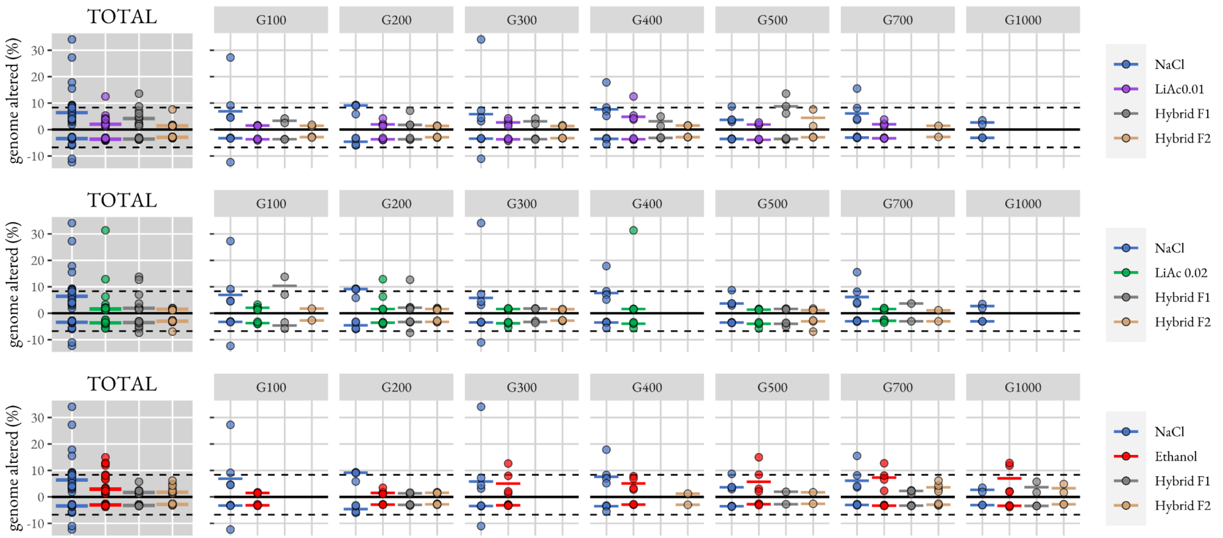


**Figure S13: The copy number variant mutational landscape of hybridization.** The proportion of the genome amplified is shown as positive numbers and the proportion deleted is shown as negative numbers. Gray plots show data averaged across all sampled timepoints (TOTAL) for the three types of hybrid crosses. Other plots show data from crosses made at 7 timepoints of increasing parental divergence (number of generations) from the initial founder populations. The solid black line indicates a pure diploid population without amplifications or deletions (% genome altered = 0). The mean proportions deleted (negative) and amplified (positive) of the two founding populations are indicated by dashed black lines.

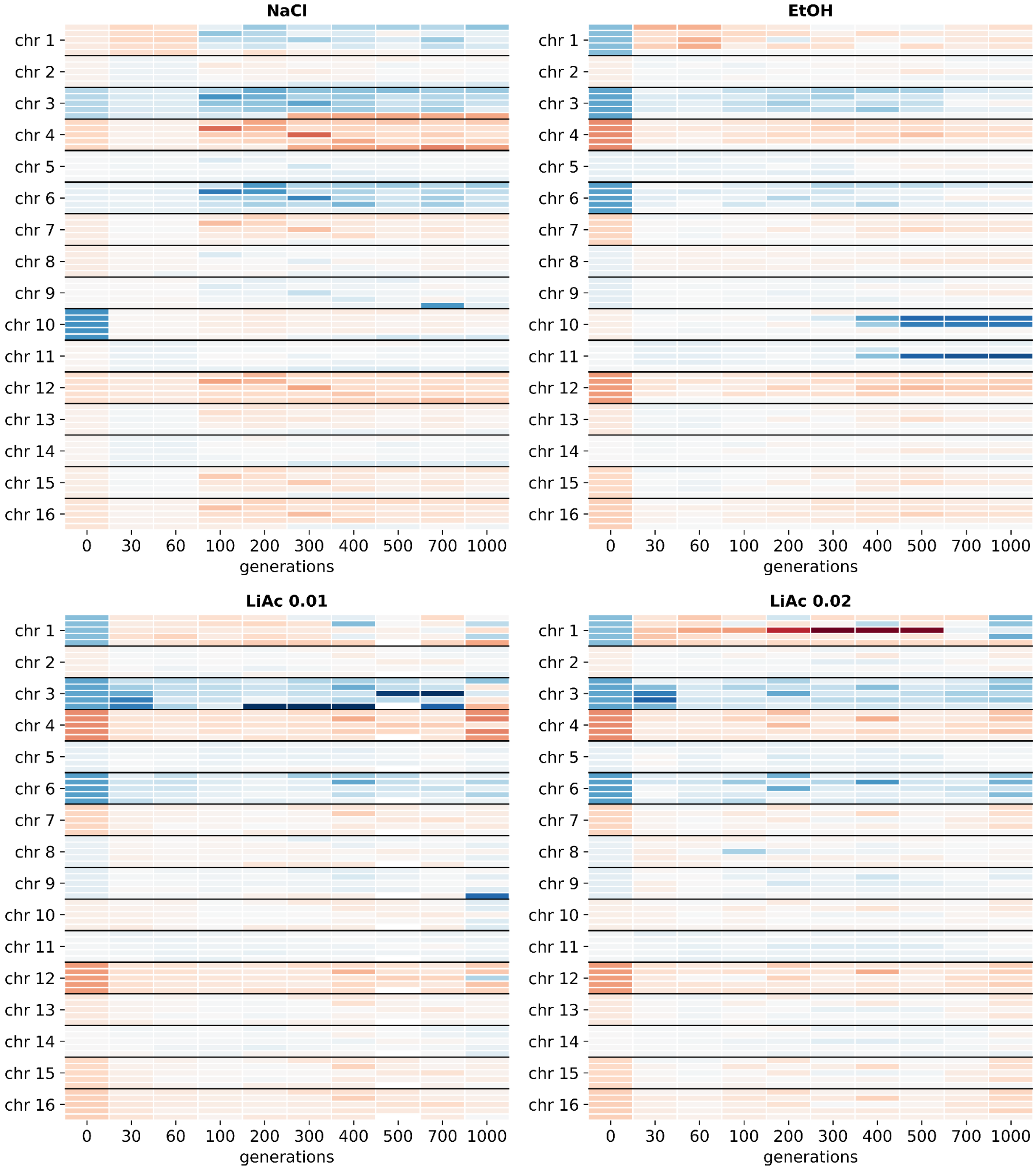


**Figure S14: Chromosome-level relative sequencing read depth.** The sequencing read depth for each chromosome relative to the genome-wide sequencing read depth for each sample. Each row indicates a replicate in each respective environment (NaCl, LiAc 0.01, LiAc 0.02 or EtOH). Blue indicates read depth greater than genome-wide average and red indicates lower than average. Sequencing timepoints not used in this study for hybrid crosses (i.e. generations 30, 60) were analyzed from (Ament-Velásquez et al. 2022).
